# Low relative air humidity and increased stomatal density independently hamper growth in young Arabidopsis

**DOI:** 10.1101/2023.10.24.563715

**Authors:** Ingmar Tulva, Kaspar Koolmeister, Hanna Hõrak

## Abstract

Stomatal pores in plant leaves mediate CO_2_ uptake for photosynthesis and water loss via transpiration. Altered stomatal density can affect plant photosynthetic capacity, water use efficiency, and growth, potentially providing either benefits or drawbacks depending on the environment. Here we explore, at different air humidity regimes, gas exchange, stomatal anatomy, and growth of Arabidopsis lines designed to combine increased stomatal density (*epf1*, *epf2*) with high stomatal sensitivity (*ht1-2*, *cyp707a1/a3*). We show that the stomatal density and sensitivity traits combine as expected: higher stomatal density increases stomatal conductance, whereas the effect is smaller in the high stomatal sensitivity mutant backgrounds than in the *epf1epf2* double mutant. Growth under low air humidity increases plant stomatal ratio with relatively more stomata allocated to the adaxial epidermis. Low relative air humidity and high stomatal density both independently impair plant growth. Higher evaporative demand did not punish increased stomatal density, nor did inherently low stomatal conductance provide any protection against low relative humidity. We propose that the detrimental effects of high stomatal density on plant growth at a young age are related with the cost of producing stomata; future experiments need to test if high stomatal densities might pay off in later life stages.

**Significance statement:** This study delves into the relationship between stomatal density, sensitivity, and environment in Arabidopsis. These findings not only enhance our comprehension of plant responses to humidity but also lay the groundwork for future studies aimed at optimising plant adaptability to varying environmental conditions.

## Introduction

Vascular plant stomata have evolved to balance CO_2_ supply for photosynthesis and water loss via transpiration (Lawson and Matthews, 2020). To perform photosynthesis efficiently, plants need a sufficient supply of CO_2_. Stomata open to allow CO_2_ to enter the leaf, which inevitably allows water vapour to escape through transpiration. While transpiration serves important functions like nutrient transport and leaf cooling, excessive transpiration can lead to water stress, especially in water-limited conditions.

Plant water loss is regulated via adjustments of stomatal conductance, determined by stomatal density and aperture. Higher stomatal density generally correlates with higher stomatal conductance (Muchow and Sinclair, 1989; Anderson and Briske, 1990; Pearce *et al*., 2006; Carlson *et al*., 2016), although it can largely be counteracted with stomatal closure (Büssis *et al*., 2006; Dow *et al*., 2014; Hepworth *et al*., 2015).

Stomatal differentiation occurs very early in leaf development, when some epidermal cells enter stomatal lineage and develop into stomata (Lee and Bergmann, 2019; Torii, 2021). During subsequent leaf expansion, stomatal density decreases (Gay and Hurd, 1975). Stomatal differentiation is governed by the sequential activation of the transcription factors SPEECHLESS (SPCH) (MacAlister *et al*., 2007), MUTE (Pillitteri *et al*., 2007), and FAMA (Ohashi-Ito and Bergmann, 2006) that is regulated by intercellular signalling mediated by epidermal patterning factors EPF2 and EPF1 (Hara *et al*., 2007; Hunt and Gray, 2009; Ohki *et al*., 2011; Zoulias *et al*., 2018). Loss-of-function mutations in the *EPF1* and *EPF2* genes lead to increased stomatal densities.

The plant hormone abscisic acid (ABA) is well known to mediate stomatal closure in response to a wide range of environmental stimuli (Seo and Koshiba, 2011; Merilo *et al*., 2013; Chater *et al*., 2015; Hsu *et al*., 2021). Arabidopsis (*Arabidopsis thaliana*) mutants with inherently high ABA levels, such as those lacking the function of some CYTOCHROME P450, FAMILY 707, SUBFAMILY A (CYP707A) proteins, exhibit increased stomatal sensitivity manifested in smaller stomatal apertures and lower stomatal conductance (Okamoto *et al*., 2006; Okamoto *et al*., 2009). CO_2_ largely regulates plant stomatal apertures via the HIGH LEAF TEMPERATURE 1 (HT1) kinase; the *ht1-2* mutants have constitutively activated CO_2_ signalling, resulting in small stomatal apertures and lower stomatal conductance (Hashimoto *et al*., 2006).

Larger stomatal densities and apertures are often accompanied by a reduced water use efficiency (Franks *et al*., 2015; Kimura *et al*., 2020). High stomatal conductance may lead to water stress, potentially impairing plant growth. Indeed, Doheny-Adams et al. (2012) demonstrate a negative correlation between stomatal density (and therefore conductance) and plant size. More recently, a series of articles have demonstrated that decreased stomatal density in Arabidopsis (Hepworth *et al*., 2015), barley (Hughes *et al*., 2017), wheat (Dunn *et al*., 2019), and rice (Caine *et al*., 2019) improves drought tolerance with no adverse effect on plant growth or carbon assimilation.

On the other hand, there is a wealth of studies that report positive relationships between stomatal density and photosynthetic assimilation or plant growth. Schlüter *et al*. (2003) demonstrated that low-light adapted Arabidopsis mutant *sdd1-1* with increased stomatal density achieved 30% higher CO_2_ assimilation rates after transfer to high light conditions than wild type. Xu and Zhou (2008) report increased net CO_2_ assimilation and even instantaneous water use efficiency with increasing stomatal density in *Leymus*. Tanaka *et al*. (2013) achieved a significant enhancement of steady-state photosynthetic capacity by overexpressing the stomata-inducing peptide STOMAGEN in Arabidopsis, and smaller biomass in a STOMAGEN-RNAi line with decreased stomatal density. Sakoda *et al*. (2020) describe 26% increase in plant biomass gain in the *epf1* mutant compared with wild-type under fluctuating light and no difference under constant light. Similarly, an increase of biomass under fluctuating, but not constant light was found in Arabidopsis mutants with large stomatal apertures (Kimura *et al*., 2020).

Combining the considerations that suggest both potential benefits and drawbacks of increased stomatal density, one might expect there to be an optimal range of stomatal conductance, subject to environmental conditions, where further increase in water loss would begin to counteract CO_2_ availability and turn detrimental to plant growth.

Experiments addressing the effects of low air humidity help to address the role of increased evaporative demand on plant growth. Low night-time relative humidity did not affect biomass production in Arabidopsis (Christman *et al*., 2009), whereas we recently found that low air humidity applied both during the day and night hampered plant growth in both wild-type plants and mutants with increased stomatal conductance due to larger stomatal apertures (Tulva *et al*., 2023).

Relatively little success has been achieved in improving plant water use efficiency and productivity, and this lack of success has been ascribed to limitations of single-gene or single-trait modification approaches (Flexas, 2016). Here, we combine mutations affecting stomatal density and sensitivity to test i) whether simultaneous manipulation of these traits leads to independent effects on stomatal conductance and could hence have potential for multi-gene applications in engineering stomatal conductance and water use efficiency, and ii) how does growth under low relative humidity affect plants with altered stomatal conductance via modification of stomatal density, stomatal sensitivity or both these traits.

In this experiment, we created Arabidopsis lines that combined mutations in *EPF2* and/or *EPF1* (leading to increased stomatal density (Hara *et al*., 2007; Hunt and Gray, 2009)) and in *HT1* or *CYP707A1* and *CYP707A3* (leading to increased stomatal sensitivity - here defined as hypersensitive stomatal closure manifested in constantly smaller stomatal apertures (Hashimoto *et al*., 2006; Okamoto *et al*., 2006; Okamoto *et al*., 2009)) and studied them for their gas exchange, stomatal anatomy, and growth under different air humidity regimes. Stomatal conductance was increased by density-altering mutations both in wild-type background and in plant lines carrying mutations that lead to smaller stomatal apertures. Moderately reduced air humidity was consistently detrimental to the size of all studied plant lines irrespective of stomatal traits. Plant lines with increased stomatal density were hampered in growth, manifesting as a negative correlation between stomatal density and plant size, whereas this reduction was not due to stomatal water loss and might instead be explained by a developmental cost.

## Materials and methods

### Plant material

The studied *Arabidopsis thaliana* mutant lines were all in Col-0 background; the single and double mutant lines were *epf1-1* (SALK_137549; Alonso *et al*., 2003; Hara *et al*., 2009), *epf2-2* (GABI_673E01; Rosso *et al*., 2003; Hunt & Gray, 2009), *epf1-1epf2-2* (Hunt and Gray, 2009), *ht1-2* (Hashimoto *et al*., 2006), and *cyp707a1-1cyp707a3* (SALK_069127 and SALK_101566, respectively; Kushiro *et al*., 2004; Okamoto *et al*., 2006); higher order mutants were obtained by crossing the single and double mutants. All plant lines used in experiments are listed in Table 1. Primers used for verifying mutant genotypes are shown in Table S1. Age-matched seed stocks were used for experiments (grown under 50% RH, 23**°**C; 10:14 h day:night until bolting, then 16:8 h day:night).

**Table 1.**
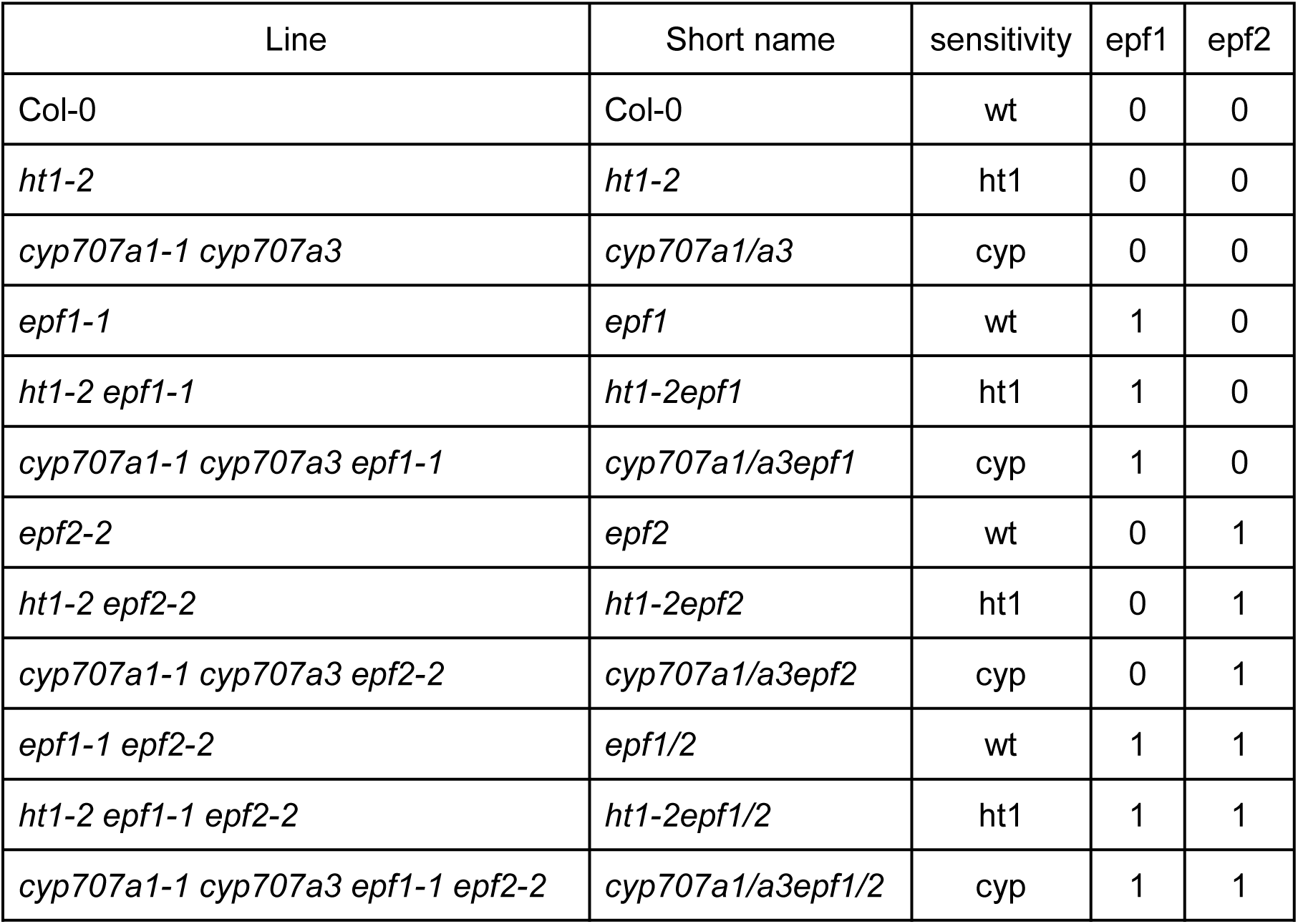
Plant lines studied, their short names used in this article, and levels of the factors used in ANOVA.

### Gas exchange experiment

Gas exchange was measured using a thermostated 8-chamber Arabidopsis gas exchange system (Plantinvent OÜ, https://www.plantinvent.com/) that allows measurement of whole-rosette leaf conductance and net assimilation rate (Kollist *et al*., 2007; Hõrak *et al*., 2017). Six plant lines were studied: Col-0, *ht1-2*, *cyp707a1/a3*, *epf1/2*, *ht1-2epf1/2*, and *cyp707a1/a3epf1/2*. The soil was a 2:1 v:v mixture of peat (OPM 025 W, Kekkilä Oy, Vantaa, Finland) and vermiculite (Vermikuliit Medium 0-4 mm, TopGreen, Vahi, Estonia). The plants were sown into pots where the soil was separated from the aboveground part by a glass, which would later form the bottom of the measuring cuvette. The plants were divided between two Percival AR-22L (Percival Scientific, IA, USA) growth cabinets, which were set to keep a 10:14 hour day:night cycle, with relative humidity (RH) of 60%:80% and air temperature of 23:19°C day:night, respectively, and daytime photosynthetic photon flux density (PPFD) of 250 μmol m^-2^ s^-1^ in both for the first 14 days, then one cabinet was switched to the low RH conditions (40%:50% day:night air humidity, otherwise identical to the control). The growth cabinets stood in a ventilated room, ensuring CO_2_ concentration of about 420 ppm around the plants. Plants were watered every few days by soaking the pots in a tray filled with shallow water to take the soil water content to ∼70-80%. Soil moisture in the dry air cabinets decreased faster than that in the control treatment; this was compensated for by more frequent watering, but no precisely defined watering regime was applied. The experiment was performed twice, for total sample size per line per growth condition of 9-15.

Gas exchange was measured in 22-27 days old plants. Measurement cuvette was thermostated to keep the air temperature at 23⁰C; incident PPFD inside the cuvette was 250 μmol m^-2^ s^-1^. Stabilised outside air with [CO_2_] about 420 ppm was used to supply air into the chambers, with only RH and temperature altered. The gas exchange system recorded one data point each minute, cycling between eight cuvettes, yielding one data point per plant every eight minutes. Plants were at first kept at ∼60% RH for 1.5 h, after which air humidity was lowered to ∼40%. Points measured between -24 to 0 minutes before humidity lowering (3-4 data points per plant) were averaged to yield steady-state 60% humidity gas exchange, and points recorded between 36 and 60 minutes after humidity lowering likewise averaged to yield 40% RH gas exchange.

The variables presented are leaf conductance (stomatal+cuticular conductance) g_l_ and net CO_2_ assimilation A_net_. Cuvette boundary layer conductance was 4.5 cm s^-1^ as measured with wet filter paper plant models during prior calibration of the system - note that this value represents both leaf sides of a typical Arabidopsis plant in this chamber, while in nature, boundary layer conductances may differ between leaf sides.

After the gas exchange measurement, the sixth leaf was collected, cut in half after weighing, and a surface imprint was taken from each half with dental silicone as in (Casson *et al*., 2009). Abaxial surface impression was collected from one half of the leaf (oranwash L, Zhermack; www.zhermack.com) and adaxial surface impression from the other (Speedex light body, Coltene/Whaledent AG; Alstätte, Switzerland). Secondary imprints were taken from the silicone with nail varnish and transferred to microscopy slides. One location (0.0675 mm^2^) from each imprint was photographed under microscope (Kern OBF 133; Kern & Sohn GmbH; www.kern-sohn.com) at 400X magnification, to be analysed for stomatal density (SD) with ImageJ software (National Institutes of Health, USA; (Schneider *et al*., 2012)).

### Growth experiment

Plants were grown in 50x29 cm trays filled with 6 cm deep soil. Water was added to the outer tray and absorbed by the soil in the inner tray through 40 3 mm holes drilled into the inner tray’s bottom.

Soil was added to the trays in two stages. First, 1200 g of 2:1 peat and vermiculite mixture (v/v; same materials as in the gas exchange measurement) was weighed into each tray. ∼50 g soil samples were taken after each 1-2 trays, dried at 70°C for two days, and weighed to determine mass based water content in the soil mixture, which was 44±1%. Based on this knowledge, 220 g of the same mixture was added to each tray, to take the total amount of estimated dry soil in a tray to 800 g, and 3 litres of water was gradually added to the soil. After the final filling, the soil in the trays was covered with black textile mulch blankets (Baltic Agro AS, Lehmja, Estonia). The mulch sheets had 32 holes (5 mm diameter, 65 mm distance between hole centres) for plants.

One plant of each Arabidopsis line to be studied was sown into a hole in the central part of the tray, randomly chosen for each tray (Figure 1). Other plants grown in those trays were not part of this study. After sowing, the trays were wrapped in cling film for 4 days to keep saturating air humidity for germination, then transferred to two Percival AR-22L growth cabinets. Conditions in the chambers were the “control” conditions as described in the gas exchange experiment. The trays were rotated and shuffled inside and between the cabinets daily.

**Figure 1.**
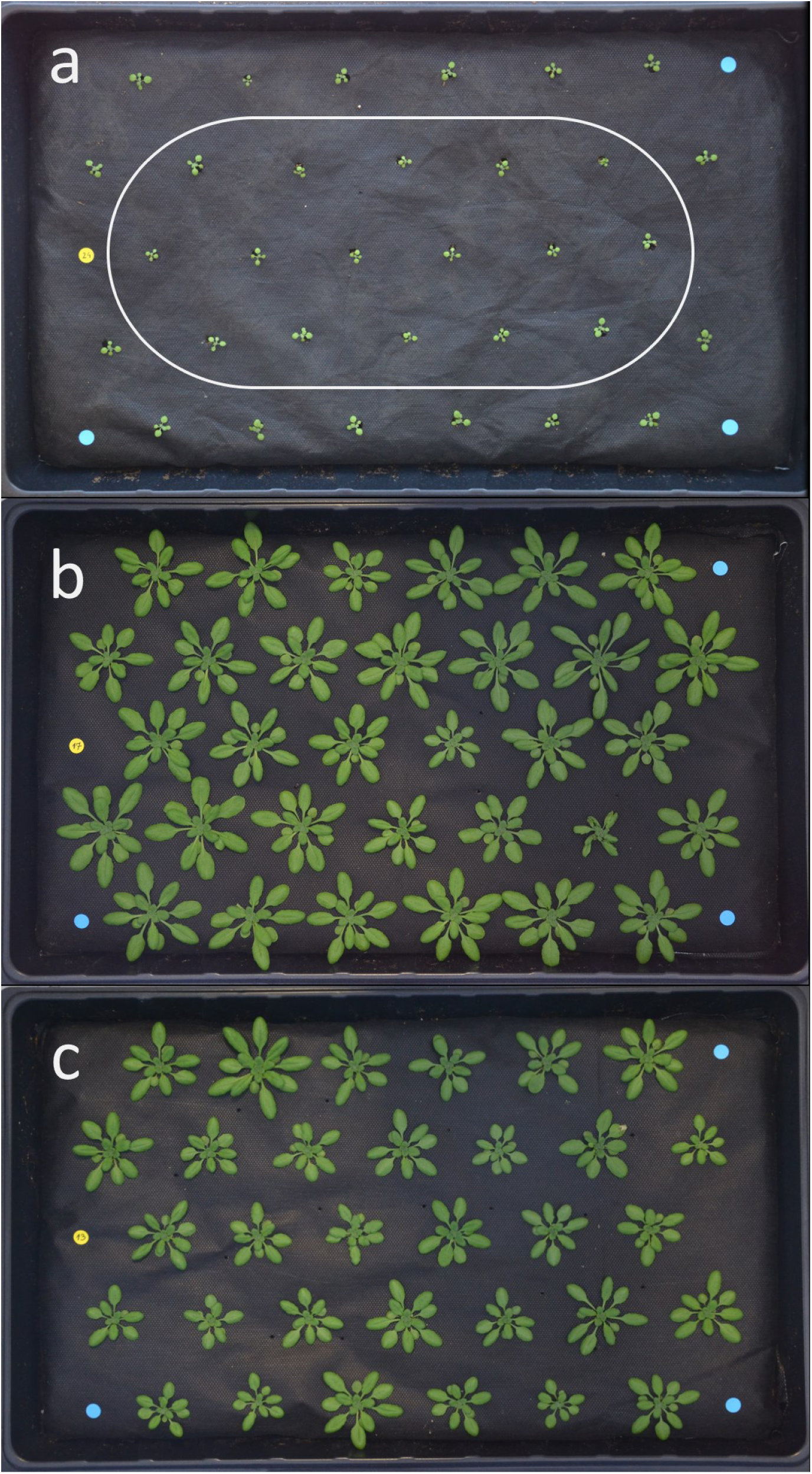
Layout of the trays in the growth experiment. (a), an example tray on Day 0 (14th day after sowing). The region surrounded by the oval included the studied plants in randomised positions, others outside were Col-0. (b), an example control tray on Day 14; (c), an example low-RH tray on Day 14.

On the 14^th^ day after sowing, one of the growth cabinets was set to the “dry” regime. From that moment onwards, the trays were rotated and shuffled around within each cabinet. Every day in the late afternoon, the trays were weighed for their gross mass to determine soil water content (SWC). When SWC in any tray was below 70%, water was added to the lower tray to take SWC to 75%. This was to compensate for the faster drying of soil in the low RH treatment, establishing a SWC pattern in each tray that would systematically fluctuate between 70% and 75% regardless of RH.

The trays were photographed daily with a Nikon D3100 camera positioned about 1 m directly above the centre of the tray. Photos were stored in Nikon RAW format, converted to Windows Bitmap with IrfanView batch conversion feature, and analysed with custom-made software for the projected area of each studied plant.

After 14 days of differential treatment, all studied plants were collected at the age of 4 weeks. The sixth leaf was separated, this leaf and the rest of the rosette were weighed for wet mass and photographed for (gently flattened) area. The remaining rosette was dried for 5 days at +60℃ and weighed for dry mass, which was then scaled for whole rosette dry mass based on the ratio of wet masses of the sixth leaf and the remaining rosette. Mass loss of the rosette (without the separated sixth leaf) during oven drying gave water content, and its dry mass divided by its harvested area gave leaf mass per area (LMA).

The sixth leaf was studied for its stomatal anatomy as described above. Additionally, from each imprint, five typical mature stomata were measured for their length (length of the whole stomatal complex); median of which was used to represent a given leaf side in stomatal size analysis. There were issues with imprint quality; only about half of the leaves could be analysed for their stomatal anatomy on both sides. Since this loss of data was purely technical and did not correlate with plant line or treatment, we regard the obtained sample as representative of the whole studied material. Multiplying SD with the area of the sixth leaf yielded the number of stomata per leaf (stomatal count, SC).

The experiment was repeated with the growth cabinets swapped, resulting in an overall sample size of 14-16 plants per treatment and genotype.

### Statistical analysis

Comparisons between measured variables were done with analysis of variance using R scripting language version 4.2.1. Type 3 sums of squares were ensured with the use of the function Anova(,type=”III”) from the library car (https://r-forge.r-project.org/projects/car/) for obtaining ANOVA results. For post-hoc tests, Tukey’s HSD for unbalanced samples was used, employing library agricolae (https://myaseen208.github.io/agricolae/). Level of significance was always set to 0.05.

For analysis of variance, studied genotypes were used as factors as follows (Table 1). In the growth experiment dataset, “epf1” and “epf2” were separate binary variables. In the gas exchange dataset, one variable “epf1/2” indicated whether both *EPF1* and *EPF2* were mutant or wild-type (no single mutants in these genes were studied). The other genes involved, *HT1*, *CYP707A1*, and *CYP707A3* (the latter two always at the same state) were encoded into a three-state variable “sensitivity” with values of “wt” (all three intact),”ht1” (*ht1-2* mutation), or “cyp707a1a3” (*cyp707a1/a3* double mutant) - note that the design lacks the combination of ht1-2 and cyp707a1a3 mutations and is thus not fully factorial. Growth RH was a two-state variable with values “control” or “dry” (60%:80% or 40%:50% day:night RH, respectively).

## Accession numbers

- *CYP707A1* – AT4G19230
- *CYP707A3* – AT5G45340
- *EPF1* – AT2G20875
- *EPF2* – AT1G34245
- *HT1 – AT1G62400*

## Results

### High SD and high stomatal sensitivity affect leaf conductance largely independently

To test whether mutations that alter stomatal density and sensitivity affect plant gas-exchange independently, we carried out experiments with mutants expected to combine high stomatal density (*epf1/2*) with small stomatal apertures (*ht1-2*, *cyp707a1/a3*). Gas exchange measurements were consistent with the expected behaviour of the studied lines: under both normal and low RH, plants deficient in both *EPF1* and *EPF2* had significantly higher leaf conductance than the corresponding lines with intact *EPF*s (Figure 2a; Figure S1; Table 2, significant main effect of epf1/2). Leaf conductance of *ht1-2* was significantly lower than wild-type under normal RH, whereas both *ht1-2* and *cyp707a1a3* had lower than wild-type g_l_ under low RH conditions. In combination with the *epf1* and *epf2* alleles these mutations expected to decrease stomatal apertures led to lower leaf conductance compared with the *epf1/2* double mutant under normal RH (Figure 2a). Low RH generally decreased leaf conductance (Table 2, significant main effect of RH), with the most prominent effect in the lines with the *epf1/2* mutations (Figure 2a). Growth conditions had little lasting effect on stomatal behaviour: only the *epf1/2* plants grown under low RH had decreased leaf conductance under low cuvette RH compared to the same genotype grown under normal RH (Figure S2).

**Figure 2.**
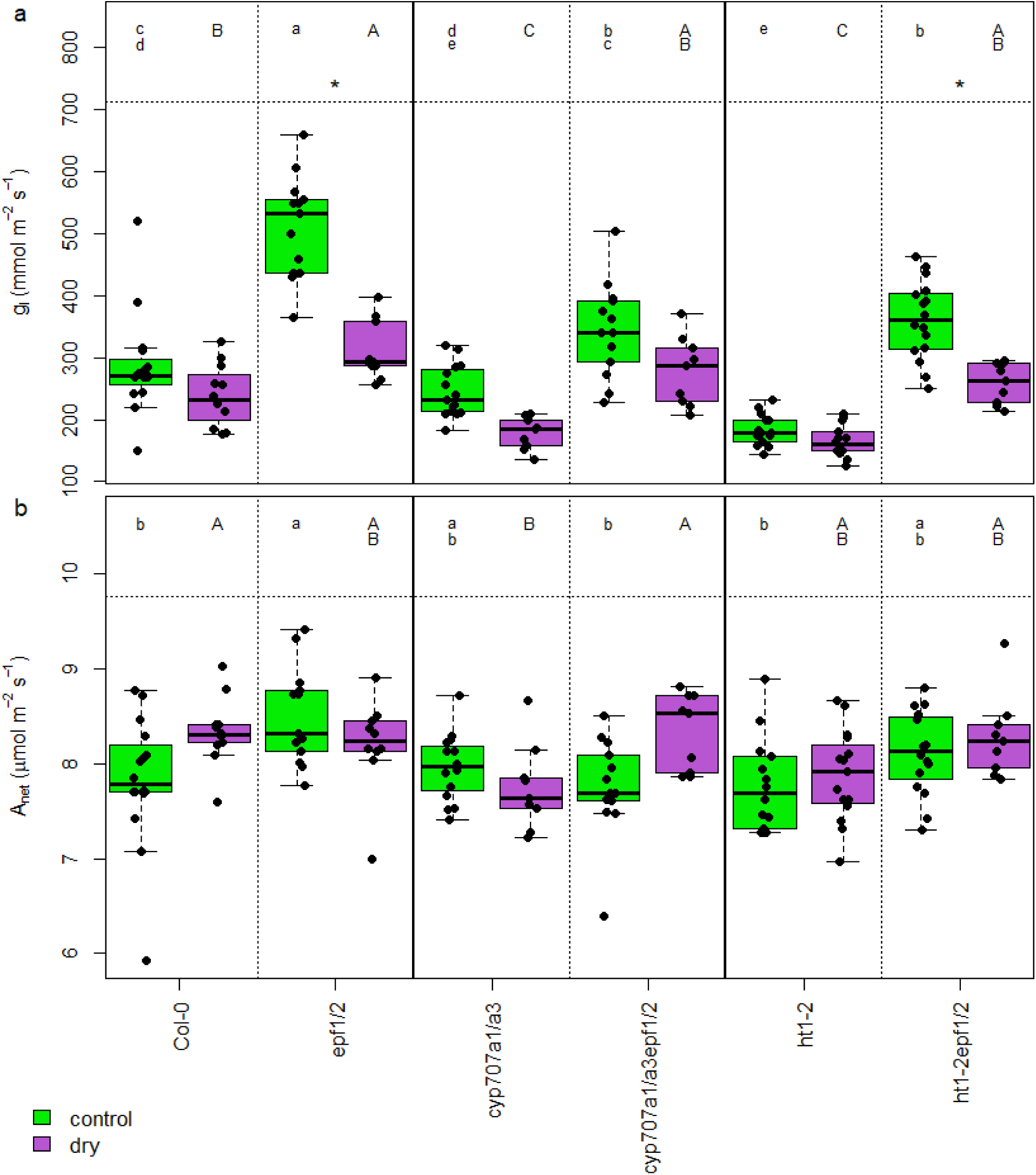
Leaf conductance to water vapour (a) and net photosynthesis (b), measured with RH in the cuvette equal to the growth RH for each plant. Green boxes, normal growth RH; magenta boxes, low RH. The boxes span from the first to the third quartile, with median indicated with the horizontal line; the whiskers span the non-outlier range. Individual plants shown as solid dots. Asterisk above a pair of boxes indicates significant difference within that genotype according to Tukey’s HSD (ANOVA involving all 24 treatment and genotype combinations). Shared lower case letters above the normal RH boxes indicate no significant difference according to Tukey HSD test (ANOVA involving only normal RH treatment); shared capital letters above the low RH boxes likewise for low RH treatment.

**Table 2.** P-values of ANOVA on steady-state gas exchange experiment. g_l_, leaf conductance; A_net_, net assimilation; WUE, intrinsic water use efficiency (=A_net_/g_l_); SD, stomatal density. Single EPF mutants were not included in this experiment; the factor epf1/2 encodes double mutant in *EPF1* and *EPF2* genes vs wild-type. RH, growth air humidity (60 % or 40 % at daytime); sensitivity encodes the state of stomatal reaction genes (*ht1-2* mutant, *cyp707a1/a3* double mutant, or wild-type). P-values < 0.05 shown in bold.

| | $g_l$<br>(growth RH) | $A_n$<br>(growth RH) | intrWUE<br>(growth RH) | SD<br>abaxial | SD<br>adaxial |
| --- | --- | --- | --- | --- | --- |
| RH | **0.00** | 0.08 | **0.00** | **0.00** | **0.00** |
| *epf1/2* | **0.00** | **0.00** | **0.00** | **0.00** | **0.00** |
| sensitivity | **0.00** | **0.02** | **0.00** | **0.00** | 0.34 |
| RH* *epf1/2* | **0.00** | 0.92 | 0.72 | 0.22 | **0.02** |
| RH*sensitivity | **0.01** | 0.93 | 0.69 | 0.23 | 0.18 |
| *epf1/2* *sensitivity | 0.11 | 0.65 | **0.00** | **0.00** | 0.66 |
| RH* *epf1/2* *sensitivity | **0.01** | **0.00** | 0.19 | 0.17 | 0.24 |

Under normal humidity, area-based net CO_2_ assimilation of the *epf1/2* mutant was higher than most other plant lines (Figure 2b). Under low RH, the *cyp707a1/a3epf1/2* line had significantly higher A_net_ than the *cyp707a1/a3* (Figure 2b). Due to decreased g_l_, the intrinsic water use efficiency (iWUE=A_net_/g_l_) generally increased under low RH (Figure 3a). Under low RH, water use efficiency was higher than in wild-type in plant lines with increased stomatal sensitivity (*ht1-2* and *cyp707a1/a3*) and about two times lower in the plant lines with the *epf1/2* double mutation than the corresponding lines with intact EPFs. Stomatal density was not affected by the *ht1-2* and *cyp707a1/a3* mutations: the *epf1/2* mutations increased both adaxial and abaxial stomatal density in all genetic backgrounds (Figure 4). Growth under low RH also increased both adaxial and abaxial stomatal density (Table 2, significant main RH effect; Figure 4), with the most prominent effect in the *epf1/2* mutant (significant RH*epf interaction in Table 2; Figure 4). Across all plant lines, the relationship between total stomatal density and leaf conductance remained positive under both growth conditions, but had a steeper slope under normal humidity (Figure 3b).

**Figure 3.**
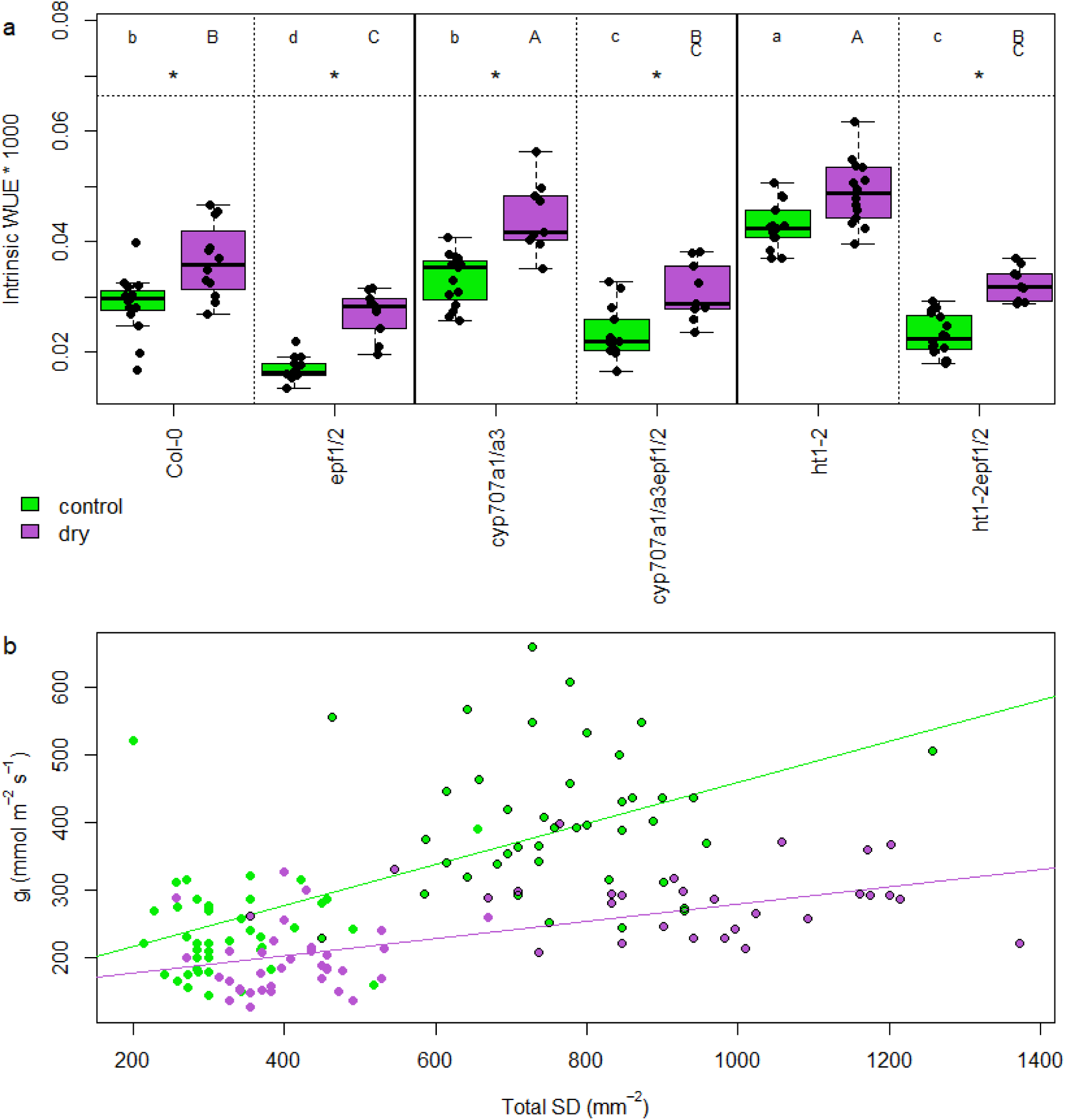
(A) Intrinsic water use efficiency in the gas exchange experiment, measured with RH in the cuvette equal to the growth RH for each plant. Green boxes, normal growth RH; magenta boxes, low RH. The boxes span from the first to the third quartile, with median indicated with the horizontal line; the whiskers span the non-outlier range. Individual plants shown as solid dots. Asterisk above a pair of boxes indicates significant difference within that genotype according to Tukey’s HSD (ANOVA involving all 24 treatment and genotype combinations). Shared lower case letters above the normal RH boxes indicate no significant difference according to Tukey HSD test (ANOVA involving only normal RH treatment); shared capital letters above the low RH boxes likewise for that treatment. (b) Relationship between stomatal density and leaf conductance at growth conditions. Each dot is one plant; the plants with the *epf1/2* double mutation are circled. Slopes of the linear regressions differ significantly from zero (p<0.05) and from each other (p<0.001).

**Figure 4.**
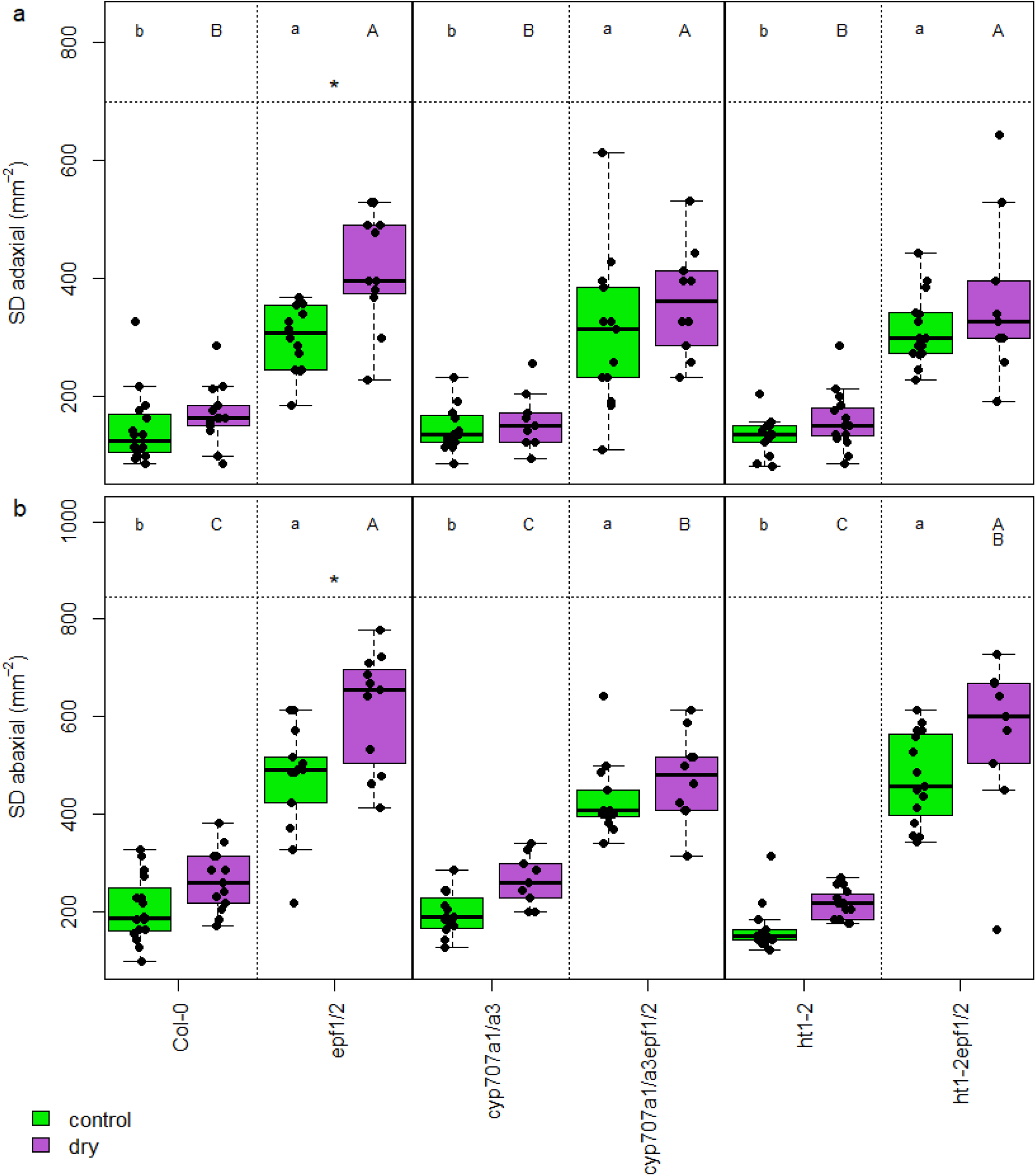
Adaxial (a) and abaxial (b) stomatal densities in the gas exchange experiment. Green boxes, normal growth RH; magenta boxes, low RH. The boxes span from the first to the third quartile, with median indicated with the horizontal line; the whiskers span the non-outlier range. Individual plants shown as solid dots. Asterisk above a pair of boxes indicates significant difference within that genotype according to Tukey’s HSD (ANOVA involving all 24 treatment and genotype combinations). Shared lower case letters above the normal RH boxes indicate no significant difference according to Tukey HSD test (ANOVA involving only normal RH treatment); shared capital letters above the low RH boxes likewise for that treatment.

### No effect of epf1 mutation on SD in plant lines with high stomatal sensitivity

To dissect the role of different stomatal density and sensitivity in plant growth under different RH environments, we carried out a growth experiment. The positive effects of the *epf1/2* and low RH on stomatal density were similar as found in the gas-exchange experiment, although low RH significantly increased only adaxial SD in the growth experiment (Figure 5; Table 3, significant RH effect). The *epf1/2* double mutants had larger stomatal density on both leaf sides than the respective lines with intact EPFs, and the *ht1-2* or *cyp707a1/a3* mutations alone did not affect stomatal density. In the Col-0 background, both *epf1* and *epf2* single mutations increased stomatal density on both leaf sides under normal RH (Figure 5), but there was a significant interaction between *epf1* mutation and stomatal sensitivity (Table 3), manifested in no increase of SD in lines combining the *cyp707a1/a3* or *ht1-2* mutations with the *epf1* single mutation (Figure 5).

**Figure 5.**
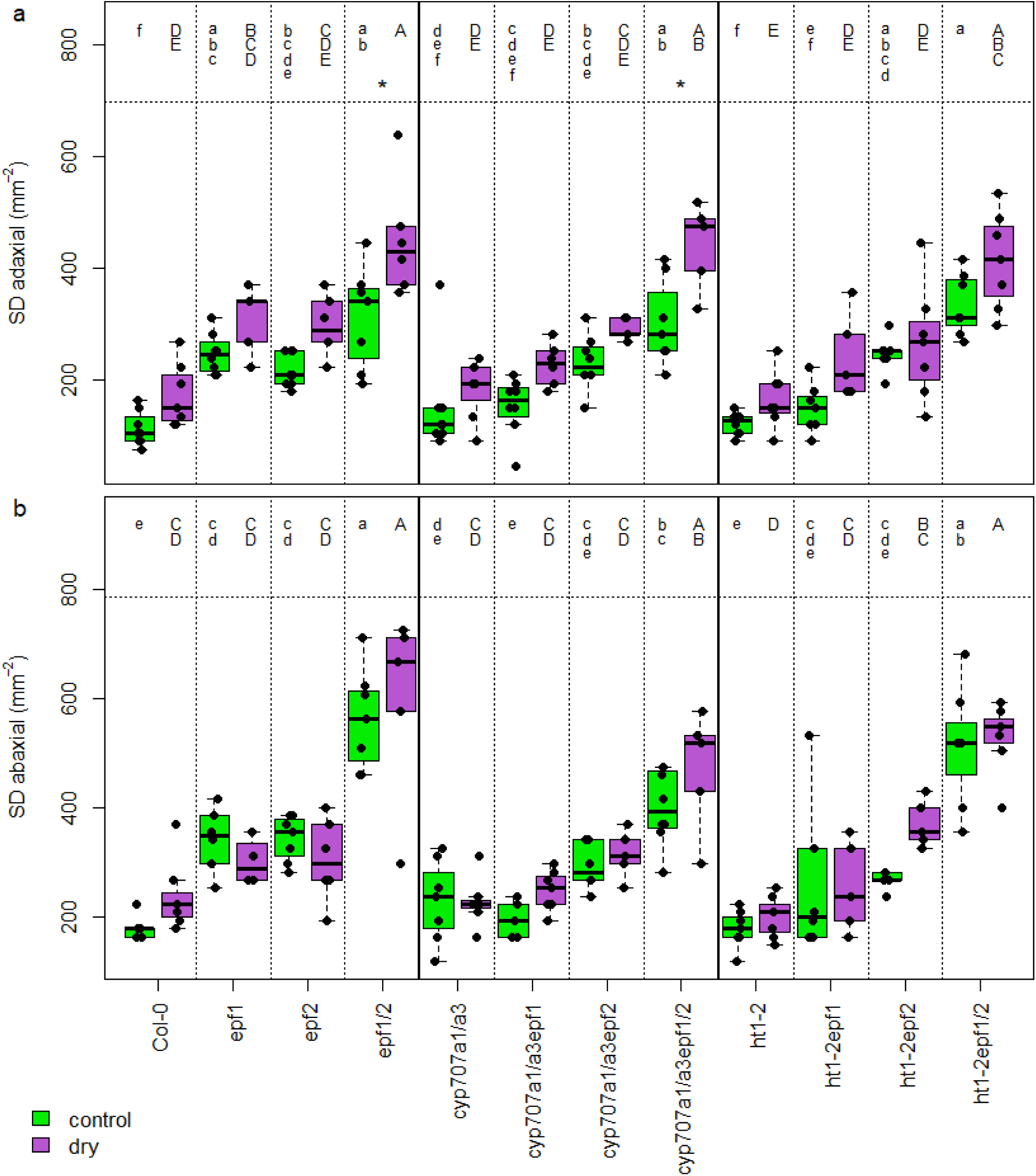
Stomatal density at the adaxial (a) and abaxial (b) leaf surface in the growth experiment. Green boxes, normal growth RH; magenta boxes, low RH. The boxes span from the first to the third quartile, with median indicated with the horizontal line; the whiskers span the non-outlier range. Individual plants shown as solid dots. Asterisk above a pair of boxes indicates significant difference within that genotype according to Tukey’s HSD (ANOVA involving all 24 treatment and genotype combinations). Shared lower case letters above the normal RH boxes indicate no significant difference according to Tukey HSD test (ANOVA involving only normal RH treatment); shared capital letters above the low RH boxes likewise for that treatment.

**Table 3.** P-values of ANOVA on the growth experiment stomatal data (collected on days 15-16 after treatment start = 29-30 after sowing). P-values < 0.05 shown in bold. SD, stomatal density; RH, growth air humidity; epf1, state of the *EPF1* gene (wild-type or mutant), epf2 likewise; sensitivity encodes the state of stomatal reaction genes (*ht1-2* mutant, *cyp707a1/a3* double mutant, or wild-type).

| Factor | SD abaxial | SD adaxial | SD sum | Stomatal count abaxial | Stomatal count adaxial | Stomatal count sum | Stomatal ratio | Stomatal length abaxial | Stomatal length adaxial |
| --- | --- | --- | --- | --- | --- | --- | --- | --- | --- |
| RH | 0.06 | **0.00** | **0.00** | **0.00** | 0.14 | **0.04** | **0.00** | **0.00** | **0.00** |
| epf1 | **0.00** | **0.00** | **0.00** | **0.00** | **0.00** | **0.00** | 0.31 | 0.77 | 0.05 |
| epf2 | **0.00** | **0.00** | **0.00** | **0.00** | **0.00** | **0.00** | 0.69 | 0.07 | 0.20 |
| sensitivity | **0.00** | 0.16 | **0.00** | **0.00** | **0.00** | **0.00** | 0.23 | 0.58 | **0.01** |
| RH*epf1 | 0.79 | **0.02** | 0.67 | 0.28 | 0.10 | 0.77 | 0.11 | 0.20 | 0.60 |
| RH*epf2 | 0.38 | 0.22 | 0.29 | 0.63 | 0.70 | 0.58 | 0.87 | 0.94 | 0.62 |
| epf1*epf2 | **0.00** | **0.02** | **0.00** | 0.27 | 0.12 | 0.82 | **0.01** | 0.92 | 0.46 |
| RH*sensitivity | 0.49 | 0.53 | 0.97 | 0.49 | 0.84 | 0.81 | 0.37 | **0.03** | 0.51 |
| epf1*sensitivity | **0.00** | **0.03** | **0.00** | **0.00** | **0.00** | **0.00** | 0.29 | 0.15 | 0.20 |
| epf2*sensitivity | 0.20 | 0.28 | 0.11 | 0.34 | 0.15 | 0.05 | 0.36 | 0.70 | 0.19 |
| RH*epf1*epf2 | 0.47 | 0.38 | 0.50 | 0.74 | 0.52 | 0.71 | 0.92 | 0.81 | 0.54 |
| RH*epf1*sensitivity | 0.19 | 0.82 | 0.41 | 0.22 | 0.66 | 0.47 | 0.22 | 0.37 | **0.03** |
| RH*epf2*sensitivity | 0.57 | 0.29 | 0.93 | 0.99 | 0.11 | 0.72 | 0.12 | 0.88 | 0.76 |
| epf1*epf2*sensitivity | 0.98 | 0.09 | 0.66 | 0.89 | **0.01** | 0.31 | 0.45 | 0.13 | 0.09 |
| RH*epf1*epf2*sensitivity | 0.16 | 0.90 | 0.40 | 0.17 | 0.93 | 0.35 | 0.58 | 0.36 | 0.91 |

Stomatal count was affected by plant genetics similarly to stomatal density: the *epf2* mutation had an overall increasing effect, while the impact of the *epf1* mutation was mediated by the state of the *cyp707a1/a3* and *ht1-2* genes (Table 3, significant epf1*sensitivity interaction). This resulted in increased stomatal numbers in the *epf1-1* line compared to the wild-type, but no significant difference in the *cyp707a1/a3epf1* or *ht1-2epf1* compared to the *cyp707a1/a3* or *ht1-2*, respectively (Figure 6). Overall, the main effect of the growth environment on the stomatal count was generally weaker than on stomatal density; only the abaxial sides of leaves had fewer stomata in plants grown under low RH (Table 3, significant RH main effect), and this effect was not significant in any particular genotype (Figure 6).

**Figure 6.**
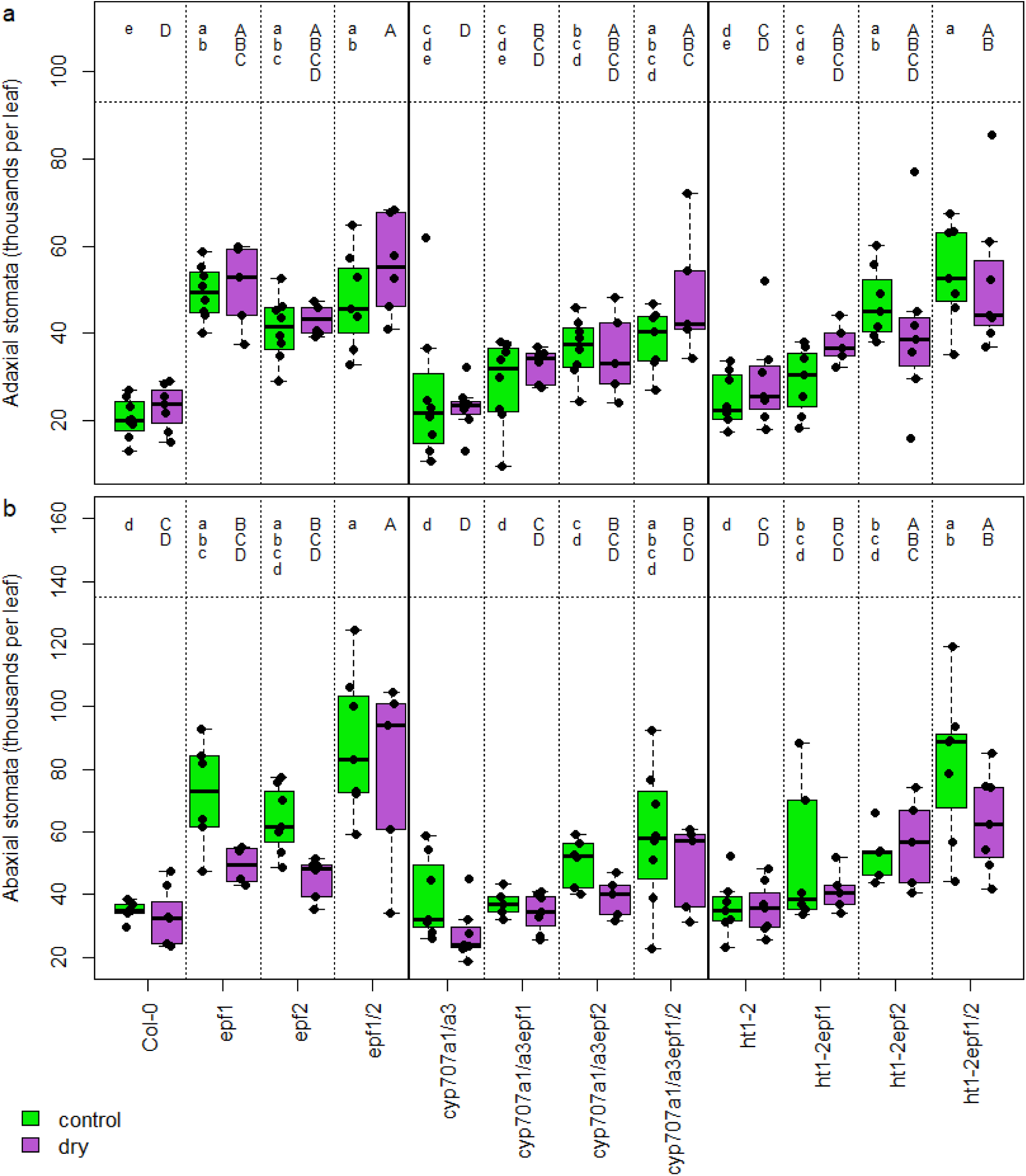
Stomatal count on the adaxial (a) and abaxial (b) leaf surface in the growth experiment. Green boxes, normal growth RH; magenta boxes, low RH. The boxes span from the first to the third quartile, with median indicated with the horizontal line; the whiskers span the non-outlier range. Individual plants shown as solid dots. Asterisk above a pair of boxes indicates significant difference within that genotype according to Tukey’s HSD (ANOVA involving all 24 treatment and genotype combinations). Shared lower case letters above the normal RH boxes indicate no significant difference according to Tukey HSD test (ANOVA involving only normal RH treatment); shared capital letters above the low RH boxes likewise for that treatment.

Growth under low RH resulted in shorter stomatal complexes on both leaf sides (Table 3, significant main effect of RH; Figure 7). Stomatal size correlated negatively but weakly with stomatal density on the adaxial side, while no significant relationship was found on the abaxial side (Figure S3).

**Figure 7.**
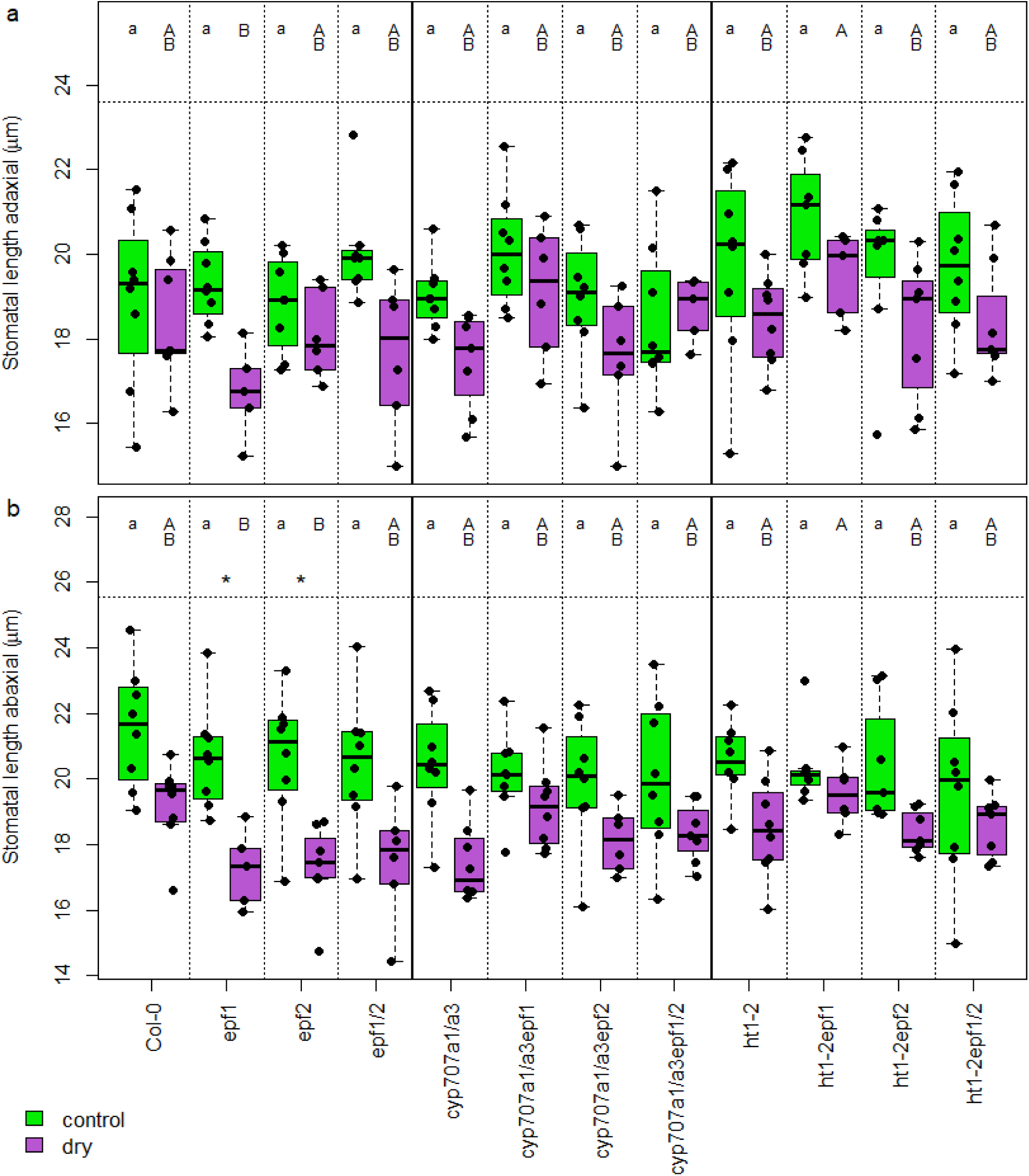
Stomatal length on the adaxial (a) and abaxial (b) leaf surface in the growth experiment. Green boxes, normal growth RH; magenta boxes, low RH. The boxes span from the first to the third quartile, with median indicated with the horizontal line; the whiskers span the non-outlier range. Individual plants shown as solid dots. Asterisk above a pair of boxes indicates significant difference within that genotype according to Tukey’s HSD (ANOVA involving all 24 treatment and genotype combinations). Shared lower case letters above the normal RH boxes indicate no significant difference according to Tukey HSD test (ANOVA involving only normal RH treatment); shared capital letters above the low RH boxes likewise for that treatment.

### Growth at low RH increases stomatal ratio

Growth under low RH slightly but consistently increased adaxial to abaxial stomatal ratio in all studied plant lines (Table 3, significant main RH effect; Figure 8; Figure S4). This effect primarily stemmed from generally increased adaxial SD under low RH, with significant increase in adaxial SD in response to low RH in the *epf1epf2* and *cyp707a1/a3epf1/2* mutants (Figure 5a; Table 3, significant RH main effect). The increase in stomatal ratio in response to low RH was strongest in plants with the *epf1* mutant allele (Figure 8).

**Figure 8.**
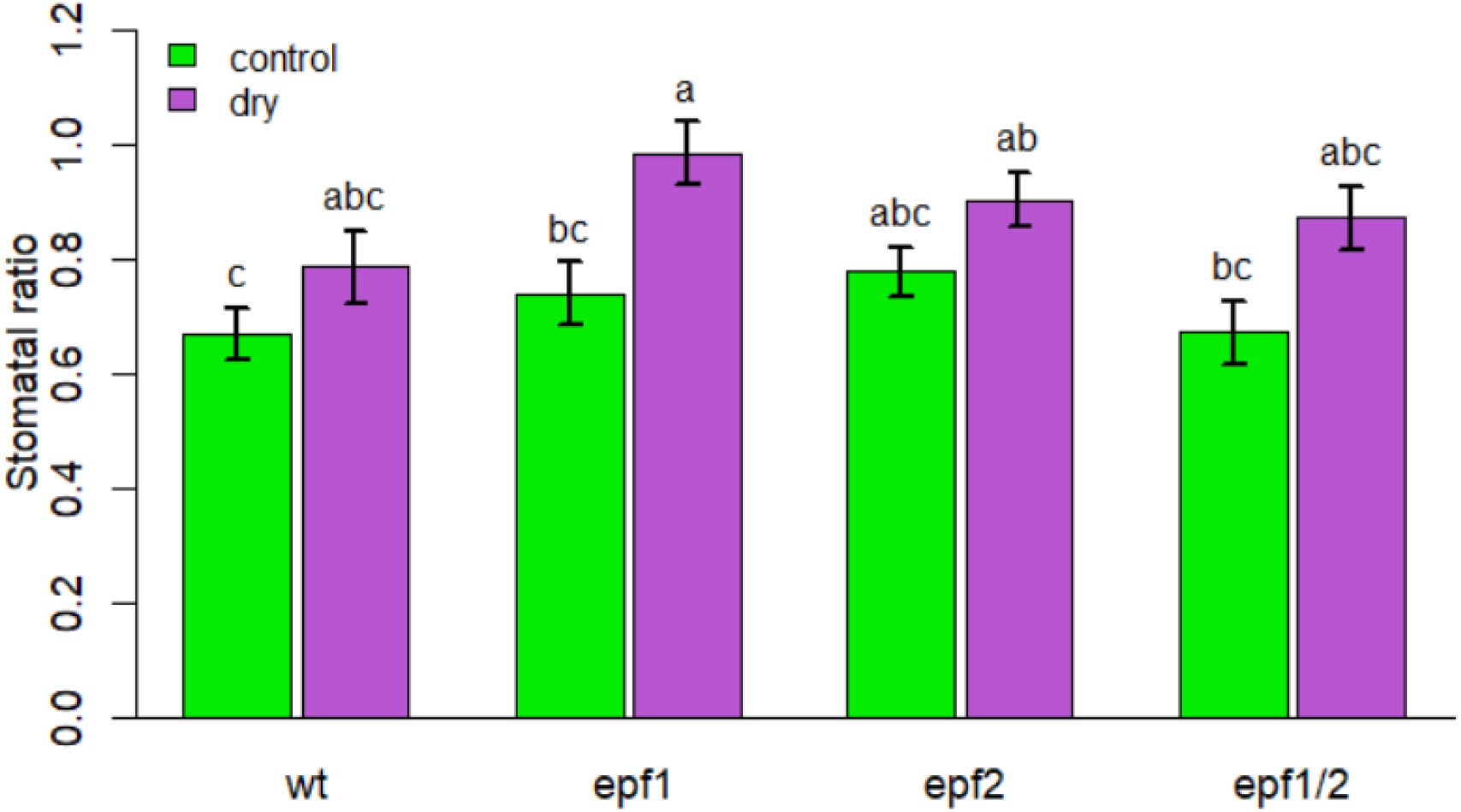
Comparison of stomatal ratio (adaxial/abaxial) in the growth experiment depending on the state of *EPF1* or *EPF2.* Stomatal sensitivity mutations, showing no significant effect on stomatal ratio (Table 3), are pooled in this data - see Figure S4 for detailed dissection. Green bars, normal growth RH; magenta bars, low RH. Shared letter(s) above the bars denote no significant difference according to Tukey HSD test (α=0.05).

### Growth at low RH and high stomatal density independently decrease plant size

Plants grown under low RH showed mild signs of xeromorphy, with slight but consistent increase in LMA and substantial decrease in water content (Figure 9; Table 4 RH main effects). A subtle increase in LMA and decrease in water content was also associated independently with both *epf1* and *epf2* mutations (Table 4, respective main effects), leading to overall positive relationship between LMA and stomatal density (Table S2, Figure S5a).

**Figure 9.**
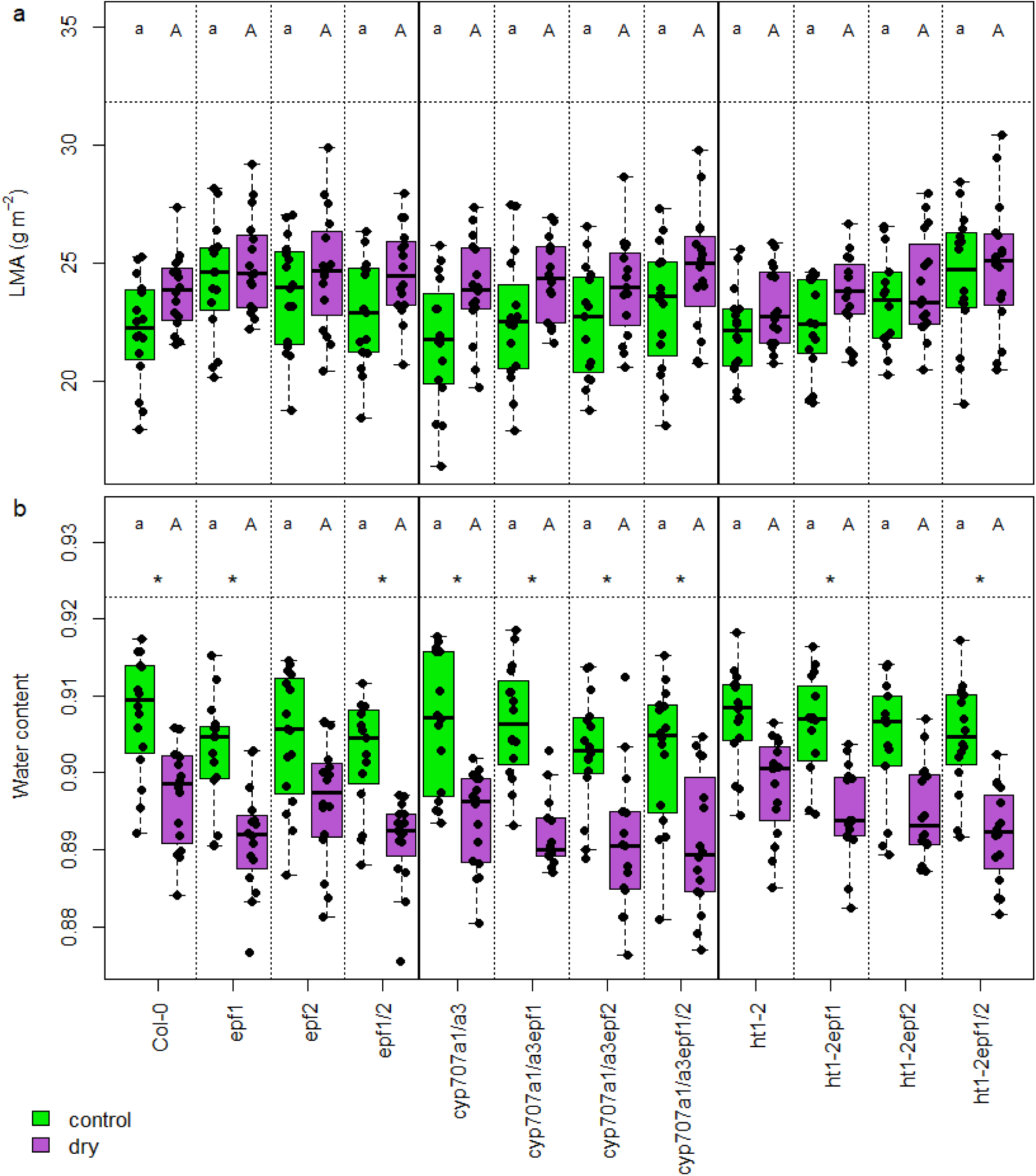
Harvest-time leaf mass / area ratio (a) and water content (b). Green boxes, normal growth RH; magenta boxes, low RH. The boxes span from the first to the third quartile, with median indicated with the horizontal line; the whiskers span the non-outlier range. Individual plants shown as solid dots. Asterisk above a pair of boxes indicates significant difference within that genotype according to Tukey’s HSD (ANOVA involving all 24 treatment and genotype combinations). Shared lower case letters above the normal RH boxes indicate no significant difference according to Tukey HSD test (ANOVA involving only normal RH treatment); shared capital letters above the low RH boxes likewise for that treatment.

**Table 4.** P-values of ANOVA on the growth experiment harvest data (collected on days 15-16 after treatment start = 29-30 after sowing), as well as projected plant areas on the day treatment started (day 0) and the last growth day (day 14; 14 and 28 days after sowing resp.). P-values < 0.05 shown in bold. LMA, leaf mass per area; RH, growth air humidity; epf1, state of the *EPF1* gene (wild-type or mutant), epf2 likewise; sensitivity encodes the state of stomatal reaction genes (*ht1-2* mutant, *cyp707a1/a3* double mutant, or wild-type).

| Factor | area<br>day 0 | area<br>day 14 | dry<br>mass | LMA | water<br>content |
| --- | --- | --- | --- | --- | --- |
| RH | 0.06 | **0.00** | **0.00** | **0.00** | **0.00** |
| epf1 | 0.96 | **0.00** | 0.13 | **0.01** | **0.00** |
| epf2 | **0.00** | **0.00** | **0.00** | **0.00** | **0.00** |
| sensitivity | 0.05 | **0.00** | **0.00** | 0.18 | 0.11 |
| RH*epf1 | 0.59 | 0.89 | 0.76 | 0.89 | 0.25 |
| RH*epf2 | 0.98 | 0.97 | 0.41 | 0.53 | 0.58 |
| epf1*epf2 | **0.01** | **0.00** | **0.01** | 0.30 | 0.41 |
| RH*sensitivity | 0.77 | **0.03** | 0.48 | 0.34 | 0.36 |
| epf1*sensitivity | 0.26 | **0.03** | 0.15 | 0.96 | 0.09 |
| epf2*sensitivity | 0.78 | 0.06 | 0.74 | 0.09 | 0.55 |
| RH*epf1*epf2 | 0.39 | 0.53 | 0.52 | 0.53 | 0.82 |
| RH*epf1*sensitivity | 0.69 | 0.16 | 0.63 | 0.98 | 0.86 |
| RH*epf2*sensitivity | 0.09 | 0.07 | 0.13 | 0.73 | 0.72 |
| epf1*epf2*sensitivity | 0.11 | 0.36 | 0.15 | **0.04** | 0.97 |
| RH*epf1*epf2*sensitivity | 0.72 | 0.42 | 0.88 | 0.61 | 0.77 |

Plant growth under low RH was significantly impaired. All plant lines showed a relatively steady decrease in dry mass at harvest time of about 25% - significantly so in the Col-0, *epf1*, *cyp707a1/a3,* and *cyp707a1/a3epf1* lines, but the trend was similar in all genotypes (Figure 10a; Table 4, significant main RH effect). The projected rosette area at the end of the experiment (28 days after sowing) decreased even more (Figure 10b), owing to the general increase in leaf mass to area ratio (Figure 9; Table 4, significant main RH effect) under low RH. Across the plant line and treatment combinations, there was a negative relationship between LMA and plant mass under low RH but not under control conditions (Figure S5b).

**Figure 10.**
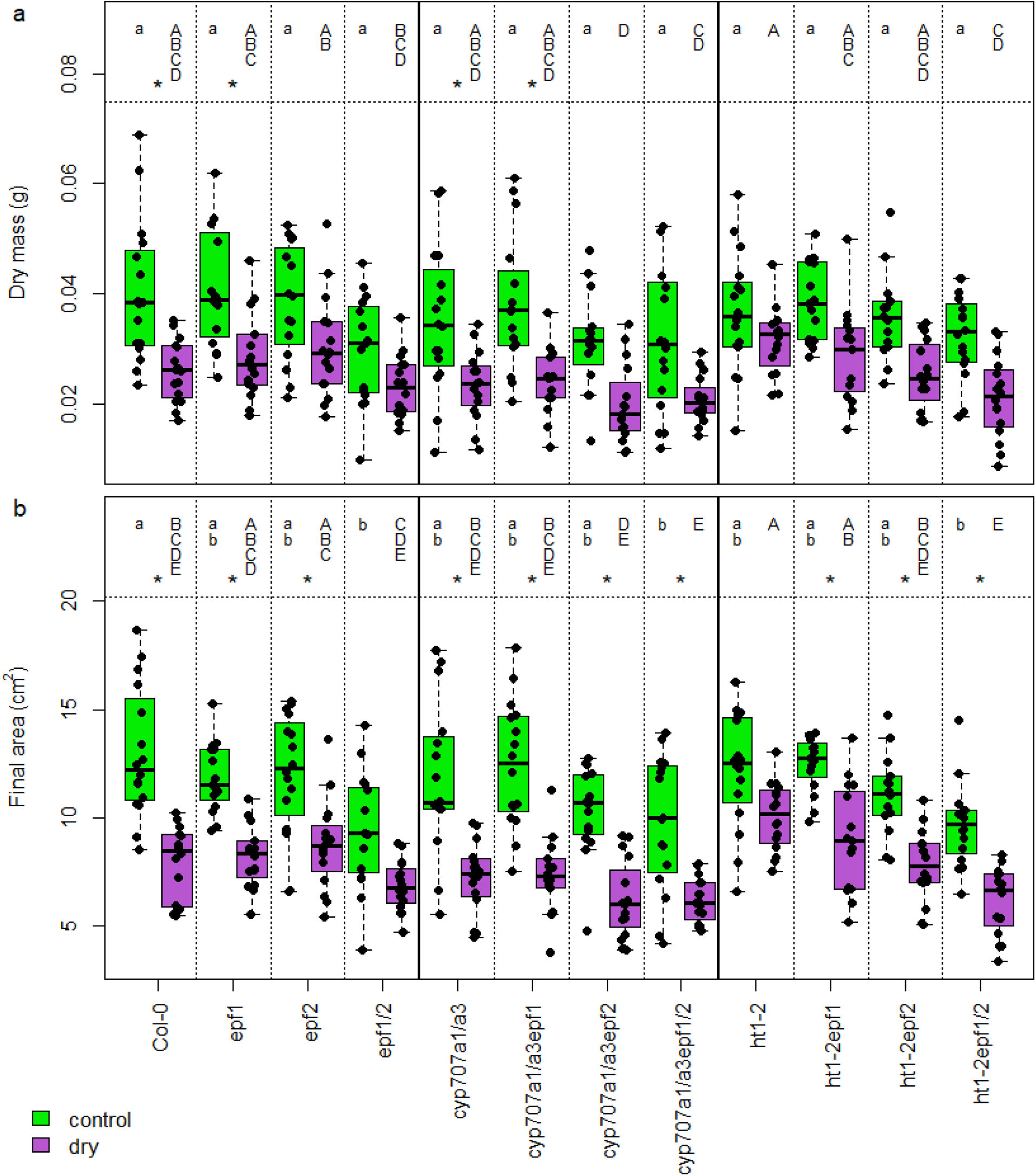
Plant above-ground dry mass (29-30 days old; a) and projected area (28 days old; b). Green boxes, normal growth RH; magenta boxes, low RH. The boxes span from the first to the third quartile, with median indicated with the horizontal line; the whiskers span the non-outlier range. Individual plants shown as solid dots. Asterisk above a pair of boxes indicates significant difference within that genotype according to Tukey’s HSD (ANOVA involving all 24 treatment and genotype combinations). Shared lower case letters above the normal RH boxes indicate no significant difference according to Tukey HSD test (ANOVA involving only normal RH treatment); shared capital letters above the low RH boxes likewise for that treatment.

Both plant area and dry mass at the end of the experiment were markedly smaller in the lines with the *epf1/2* double mutation; in case of mass, this was due to the main effect of the *epf2* mutation and there was no main effect of *epf1* (Figure 10a; Table 4), while the final area was affected by both *epf1* and *epf2* mutations (Table 4). At the onset of the RH treatment 14 days after sowing, the plant lines with the *epf2* mutations had slightly smaller projected area, a small enough effect to only appear in the pooled data (Table 4, epf2 main effect; Figure S6). During the differential treatment in weeks 3 and 4, growth under low RH resulted in a smaller relative growth rate in all plant lines, with no significant difference associated with any genetic features (Table S3, Figure S7).

As a result of the size reduction associated with the *epf1* and *epf2* mutations, particularly that of *epf1/2* double mutants, there was a significant negative correlation between total stomatal density and plant size (Figure 11a, Table S2). The slope of this relationship remained the same under both growth humidities, with the normal RH-grown plants consistently around 3.5 cm^2^ larger in their projected area, or 10 mg heavier in their dry mass. Whereas the stomatal count and plant size relationship had a similar downward trend, it was not statistically significant (Figure 11b; Table S2). Notably, the enhanced photosynthesis of *epf1/2* mutants (Figure 2b, Table 2) and overall strong correlation between stomatal density and net photosynthesis (Table S2) did not translate into improved growth but instead led to a strongly negative correlation between net photosynthesis and plant size (Table S2).

**Figure 11.**
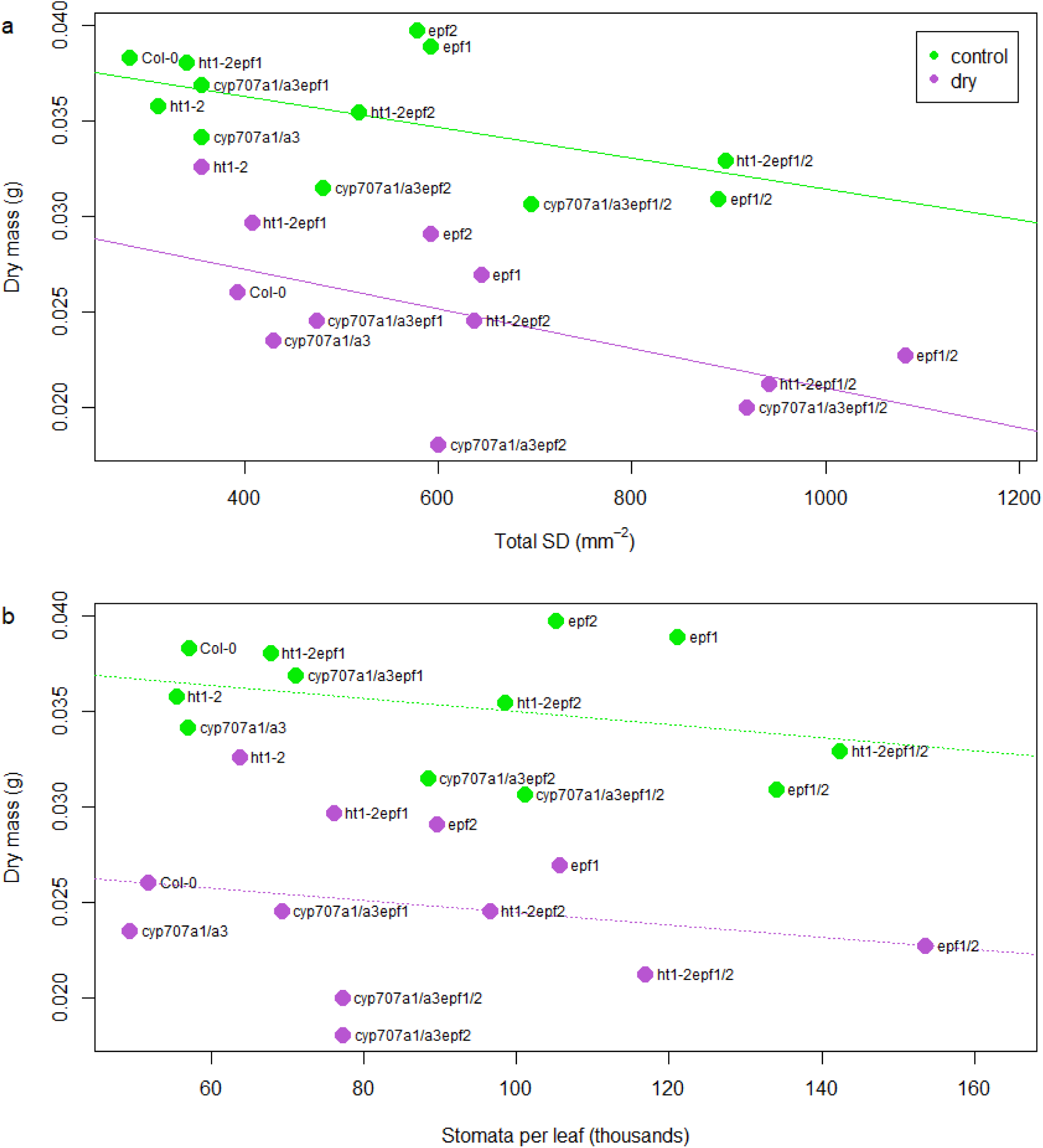
Dependence of above-ground dry mass (29-30 days old) on total stomatal density of leaf 6 (sum of both leaf sides; a) or estimated total number of stomata on leaf 6 (b). Green markers, normal growth RH; magenta markers, low RH. Each dot indicates the median values (in both dimensions) of the shown plant line in given growth conditions. The slopes of the linear regression lines in (a), themselves all significantly different from zero at P<0.01, do not differ significantly from each other between growth conditions. The slopes in (b) do not differ significantly from 0.

## Discussion

Both altered stomatal density and sensitivity impact plant water use efficiency and productivity (Doheny-Adams *et al*., 2012; Franks *et al*., 2015; Kimura *et al*., 2020; Sakoda *et al*., 2020), but the potential interaction of signalling pathways governing these traits and their combined effects on plant stomatal conductance, water use efficiency, and growth have not been addressed. Here we combined mutations in *EPF1* and *EPF2* that affect stomatal density and in *HT1* and *CYP707A1/A3* that affect stomatal sensitivity under normal and low RH conditions to address these questions. While the studied mutations that affect stomatal density and sensitivity mostly acted independently (Figures 2, 4, 5, Tables 2, 3), mutations in the *EPF1* gene stood out, as in the high stomatal sensitivity backgrounds *epf1* alone did not increase stomatal density (Figure 5, Table 3). EPF2 can partially replace EPF1 function but not vice versa (Hara *et al*., 2009); it is possible that the high stomatal sensitivity background stimulates *EPF2* expression or activity in the *epf1* single mutants, leading to normal stomatal densities. It has been proposed that a combination of multiple traits could be necessary for engineering more water use efficient and productive crops (Flexas, 2016). Our results suggest that combining traits that affect stomatal density and sensitivity could be one approach, as these traits affect stomatal conductance independently. Given the negative relationship between stomatal density and plant growth (Figure 11a), optimisation of stomatal density and sensitivity trait combinations is needed for practical applications.

Our experiments showed an increase in stomatal ratio in response to low air humidity (Figure 8, Table 3). Allocating more stomata to the adaxial epidermis under dry air conditions may seem counterintuitive, as excessive water loss via adaxial stomata has been proposed as one of the costs of adaxial stomata in plants (Buckley *et al*., 2015; Drake *et al*., 2019). Yet, a similar increase in stomatal ratio under dry air was found also in several cotton varieties (Devi and Reddy, 2018), whereas amphistomatous plant species are common in dry habitats (Wood, 1934; Muir, 2015; Bucher *et al*., 2017) and tend to have faster stomatal responses than hypostomatous plants (Haworth *et al*., 2018). It is possible that high stomatal ratios allow plants to make better use of favourable conditions when they arise, by increasing CO_2_ uptake capacity (Mott *et al*., 1982; Muir, 2018; Xiong ^a^nd Flexas, 2020^)^. At the same time, densely packed stomata are typically smaller and faster than sparse stomata (Drake *et al*., 2013), ensuring fast and efficient restriction of water loss via speedy stomatal closure upon water stress. In line with this, we found smaller stomata with potential for faster stomatal responses in plants grown under low RH (Table 3, Figure 7). In our experiments, the negative relationship between stomatal density and size that is typically found in plants (Franks and Beerling, 2009; Caine *et al*., 2023) was weakly present only in the adaxial leaf surface that had lower stomatal density than abaxial leaf surface, where no such relationship was found (Figure S3, Figure 6). These data suggest that the often observed negative relationship between stomatal density and size is not due to physical limitations of packing stomata into the epidermis but may have other underlying causes.

How plants adjust stomatal ratio in response to changes in environmental conditions is not clear. The pronounced increase in stomatal ratio under low RH in the *epf1* single mutant backgrounds suggests that EPF1 as a negative regulator of stomatal development preferentially suppresses stomatal production in the adaxial epidermis in response to a decrease in relative air humidity (Figure 8). EPF2 peptide acts on an earlier step in the stomatal differentiation pathway compared with EPF1; it is possible that its absence in the *epf1epf2* double mutant suppresses the effects of *epf1* mutation, leading to no significant increase in stomatal ratio under low RH in the *epf1epf2* double mutant (Figure 8). On the other hand, the *epf1* and *epf2* mutations are additive (Figure 5), indicating that they can largely replace the function of each other. The leaf-side specific behaviour of EPFs and other stomatal regulators has not been addressed, but given that silencing of *SPCH* expression or overexpression of a *SPCH* repressor led to preferential loss of abaxial stomata (Dow *et al*., 2014; Qi *et al*., 2019), it is likely that signalling events that govern adaxial and abaxial stomatal formation are regulated independently and at least partly via different mechanisms. The latter hypothesis is also supported by our recent study of adaxial and abaxial stomatal development patterns in plants deficient in known stomatal regulators (Jalakas *et al*., 2024). Indeed, across-species studies indicate independent evolution of adaxial and abaxial stomatal densities with a greater flexibility of adaxial SD (Muir *et al*., 2023), in line with more prominent increases in adaxial SD in response to low air humidity (Figure 5). Further studies of stomatal ratio and its responses to changes in environment in different mutants of stomatal patterning, environmental and hormonal signalling pathways are needed to understand how stomatal ratio adjustments occur in plant development.

In our experiments we found that relative air humidity and high stomatal density independently impair plant growth (Table 4, Figure 10, Figure 11). We anticipated a larger negative effect of low RH in plant lines with increased stomatal conductance that we presumed would suffer from water stress. It is well known that drought induces preferential resource allocation into roots, leading to reduced above-ground growth (Eziz *et al*., 2017) and it is possible that dry air would trigger a similar outcome. In our experiment, low RH was uniformly detrimental to all genotypes, despite compensation of high stomatal conductance caused by high stomatal density of the *epf1epf2* mutations by the increased stomatal sensitivity mutations *ht1-2* and *cyp707a1/a3* (Figures 2, 4, 5). It is possible that efficient stomatal closure under low RH compensated for increased stomatal densities in our experiment (Figure 3b). Indeed, mutants with high stomatal density can efficiently close stomata in response to changes in environmental conditions (Dow *et al*., 2014; Hepworth *et al*., 2015). However, in our recent study, growth under low RH was not more impaired in the *ost1* mutant with large stomatal conductance than in wild-type plants (Tulva *et al*., 2023). As the *ost1* mutant is deficient in stomatal closure in response to low humidity, the lack of stronger growth impairment than in wild-type plants cannot be explained by stomatal closure in response to low RH. Thus, hampered plant growth under low RH seems to occur independently of stomatal traits.

Plants with high stomatal densities were stunted in growth irrespective of conditions (Figures 10, 11, Table 4), indicating that excess stomatal development impairs plant growth during the first 4 weeks in Arabidopsis. Similar impairments have been found before in mutants with high SD (Doheny-Adams *et al*., 2012; Lawson *et al*., 2014), although other studies have reported increased photosynthesis, carbon gain or biomass production in high stomatal density mutants (Schlüter *et al*., 2003; Tanaka *et al*., 2013; Sakoda *et al*., 2020). In this study, we saw that higher stomatal density was indeed associated with increased net photosynthesis across plant lines, but resulted in reduced above-ground growth nonetheless (Table S2), indicating stronger investment of photosynthetic products into processes other than leaf growth in plant lines with higher stomatal density. Hepworth *et al*. (2016) showed a strong positive correlation between stomatal density and root growth, possibly associated with increased water demand due to high stomatal conductance. An increased investment into root production in early development in high stomatal density mutants could potentially lead to delayed above-ground growth. This mechanism, however, is not easy to reconcile with the absence of similar growth retardation in the high stomatal conductance *ost1-3* mutant in Tulva *et al*. (2023).

The negative relationship between stomatal density and plant growth may be explained by increased porosity due to higher numbers of substomatal cavities and decreased amounts of photosynthetic tissue in high density mutants. Dow et al. (2017) showed that mesophyll porosity increases together with stomatal density in Arabidopsis lines; similar coupling between stomatal and mesophyll anatomy was shown in wheat (Lundgren *et al*., 2019). Given the negative relationship between stomatal density and plant growth, but not overall leaf stomatal numbers and plant growth (Figure 11), it is possible that the proportion of leaf tissue dedicated to non-stomatal structures that is larger in plant lines with lower stomatal density contributes to their improved growth. Alternatively, the lack of significant correlation between stomatal count and plant size (Figure 11b) may reflect different expansion growth due to differences in energy allocation into cell divisions, leading to a negative relationship between stomatal density and plant size (Figure 11a). Leaf construction cost is strongly positively related with LMA (Mizokami *et al*., 2022), hence leaves with higher LMA values should be more expensive for the plant to build. Indeed, we found a positive relationship between LMA and stomatal density (Figure S5a), potentially suggesting larger construction costs of leaves with high stomatal densities. At the same time, LMA was negatively related with dry mass only under low RH but not under normal conditions (Figure S5b), suggesting that LMA as a proxy for leaf construction costs alone does not explain the reduced growth of plants with higher stomatal numbers irrespective of growth humidity (Figure 11a). More detailed analyses of leaf anatomy, including mesophyll characteristics, are needed to understand the mechanism of growth reduction in high stomatal density mutants.

It is worth noting that studies that find adverse effects of high stomatal density, e.g. Doheny-Adams et al. (2012) and this study, work with constant artificial lighting, as is common in indoor growth chambers. On the other hand, studies reporting positive relationship between plant growth and stomatal density or conductance often find such effects under fluctuating light conditions (Kimura *et al*., 2020; Sakoda *et al*., 2020), where the growth improvement probably results from immediately high stomatal conductance upon step increase in light availability, thereby reducing resistance to CO_2_.

The formation of stomata has been associated with developmental and metabolic costs, whereas cell division costs have been considered larger than cell expansion and differentiation costs (Dow *et al*., 2014; Franks *et al*., 2015; Bowers, 2018). It is possible that a stoma, as a derivative of a protodermal cell through multiple divisions and differentiations, has a greater construction cost associated with it than pavement cells. Indeed, in this study, the high-SD associated size differences were already present in day 0 when stomata must have been differentiated, and there was no difference in relative growth rate throughout 3rd and 4th growth week. The energy-consuming process of stomatal formation might lead to growth impairment in high stomatal density mutants at a young age, whereas this deficiency may well be compensated by leaf expansion in later growth stages. Then, high stomatal numbers that potentially mediate increased CO_2_ uptake may pay off in improved photosynthesis and biomass production and/or yield. For example, high stomatal conductance in Arabidopsis cauline leaves was associated with higher seed yield, similar to flag leaves in rice (Ding *et al*., 2023). Further studies addressing the effects of altered stomatal densities on plant growth, biomass production and yield in later developmental stages are needed to assess the potential benefits of increased stomatal density.

## Acknowledgements

The work to generate the mutants used in this study was initiated in the lab of prof. Julie Gray; we are grateful for her support and for the seeds of *epf1epf2* mutants. We also thank Dr. Luke Fountain for help with genotyping in the early stages of the project, Dr. Ebe Merilo for comments on the manuscript, and Mikk Välbe for keeping the lab running. This work was supported by the Estonian Research Council grants PSG404 and MOBERC104 and conducted using the Plant Biology Infrastructure TAIM funded by the Estonian Research Council (TT5).

