## Supplementary Figures for "Low relative air humidity and increased stomatal density independently hamper growth in young Arabidopsis"

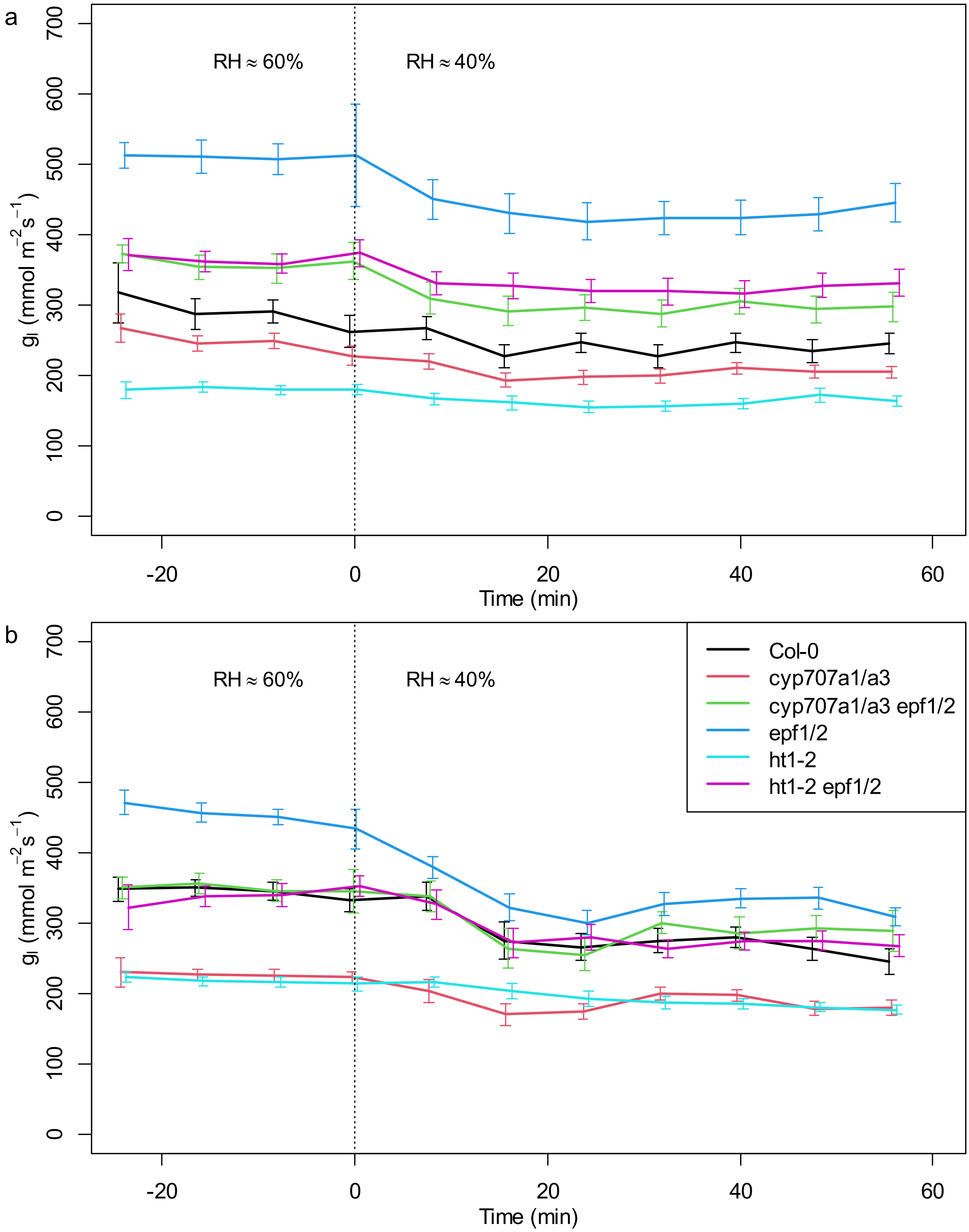


**Figure S1**. Reaction of leaf conductance to step decrease (at time 0) in cuvette air humidity. a, plants grown in control conditions (daytime RH=60%); b, plants grown in dry air (daytime RH=40%). Each marker denotes the mean±SE of an eight-minute period centred around the given marker's x position, with time points from the first four minutes after the RH switch omitted for clarity due to unstable chamber conditions and variable-strength wrong-way response. The goal of this study was to gain steady-state values for both humidity conditions, and the precise kinetics have a strong random component due to different plants having been measured at different time shifts relative to RH change.

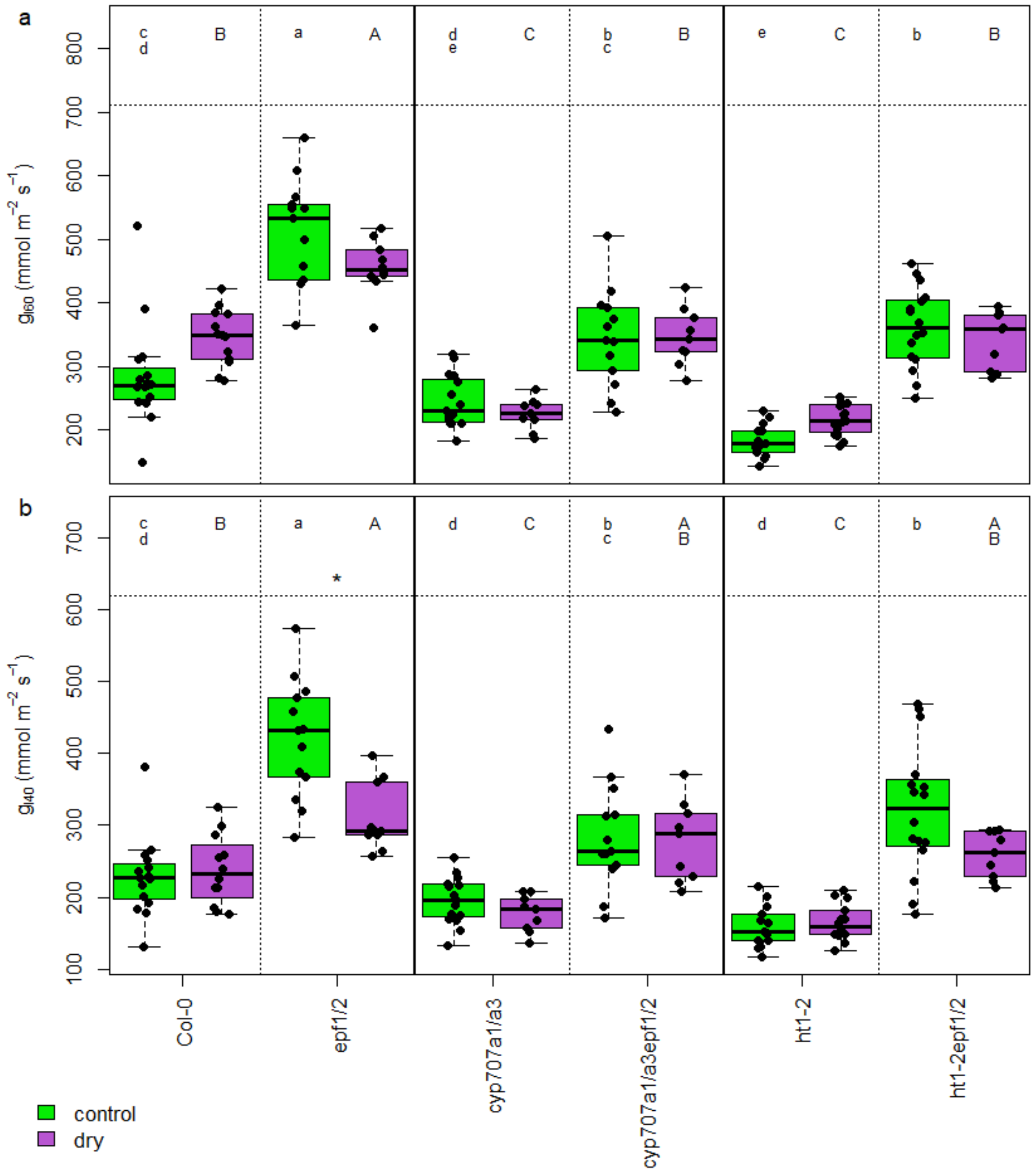


**Figure S2**. Leaf conductance at (a) 60% cuvette RH and (b) 40% cuvette RH. Green boxes, normal growth RH; magenta boxes, low RH. The boxes span from the first to the third quartile, with median indicated with the horizontal line; the whiskers span the non-outlier range. Individual plants shown as solid dots. Asterisk above a pair of boxes indicates significant difference within that genotype according to Tukey's HSD (ANOVA involving all 24 treatment and genotype combinations). Shared lower case letters above the normal RH boxes indicate no significant difference according to Tukey HSD test (ANOVA involving only normal RH treatment); shared capital letters above the low RH boxes likewise for low RH treatment.

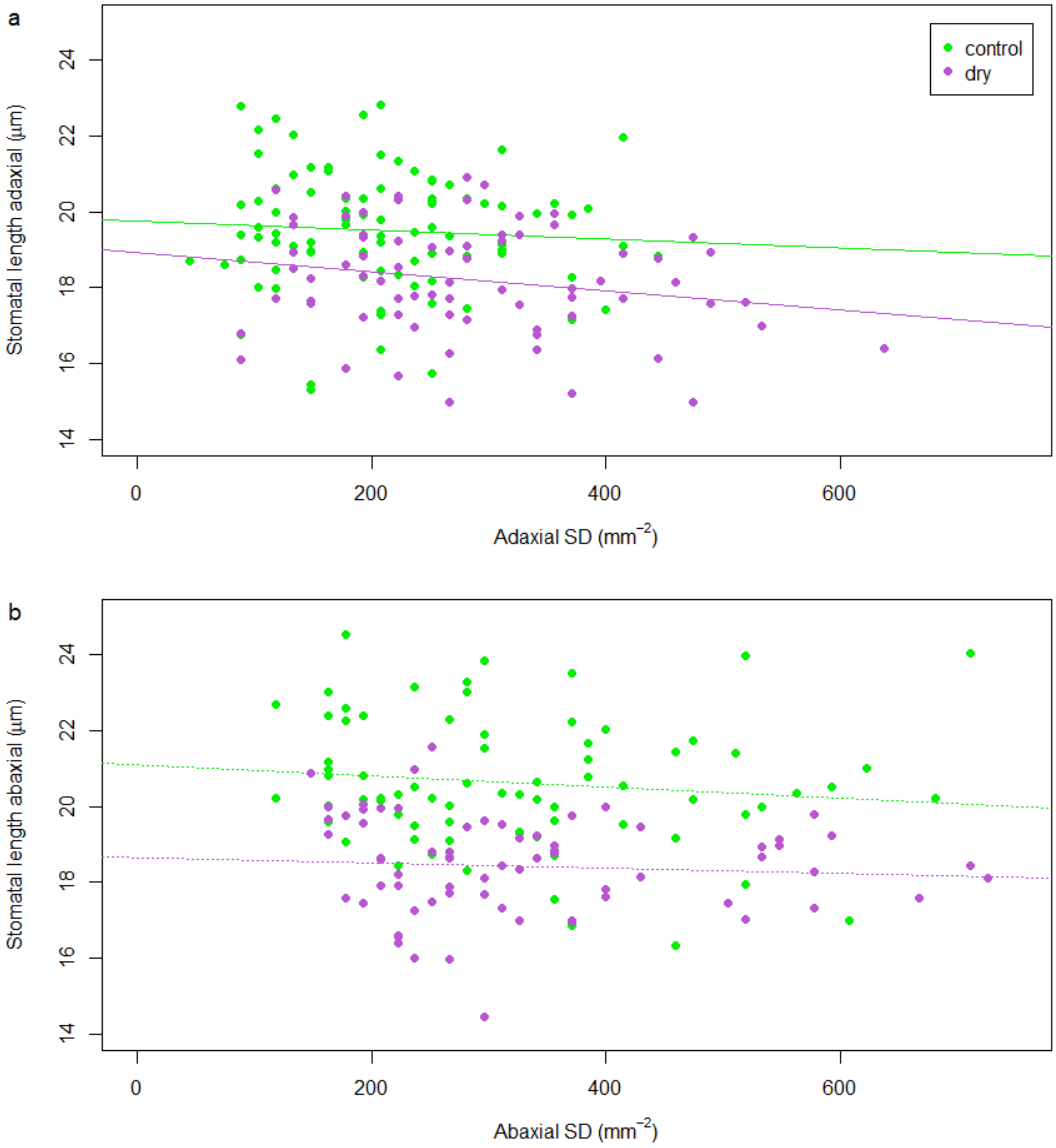


**Figure S3.** Relationship between stomatal density and stomatal length in (a) adaxial and (b) abaxial leaf side. Each marker denotes data on one leaf: density counted from one microphotograph, stomatal length measured as the median length of five mature stomata from that photo. Green markers, plants grown in control conditions; magenta markers, plants grown in reduced air humidity. In the adaxial side, the overall density vs length relationship slope differed significantly from zero ($P<0.05$) while the slopes at different growth conditions did not differ from each other; in the abaxial side, the density vs length relationship was not statistically significant.

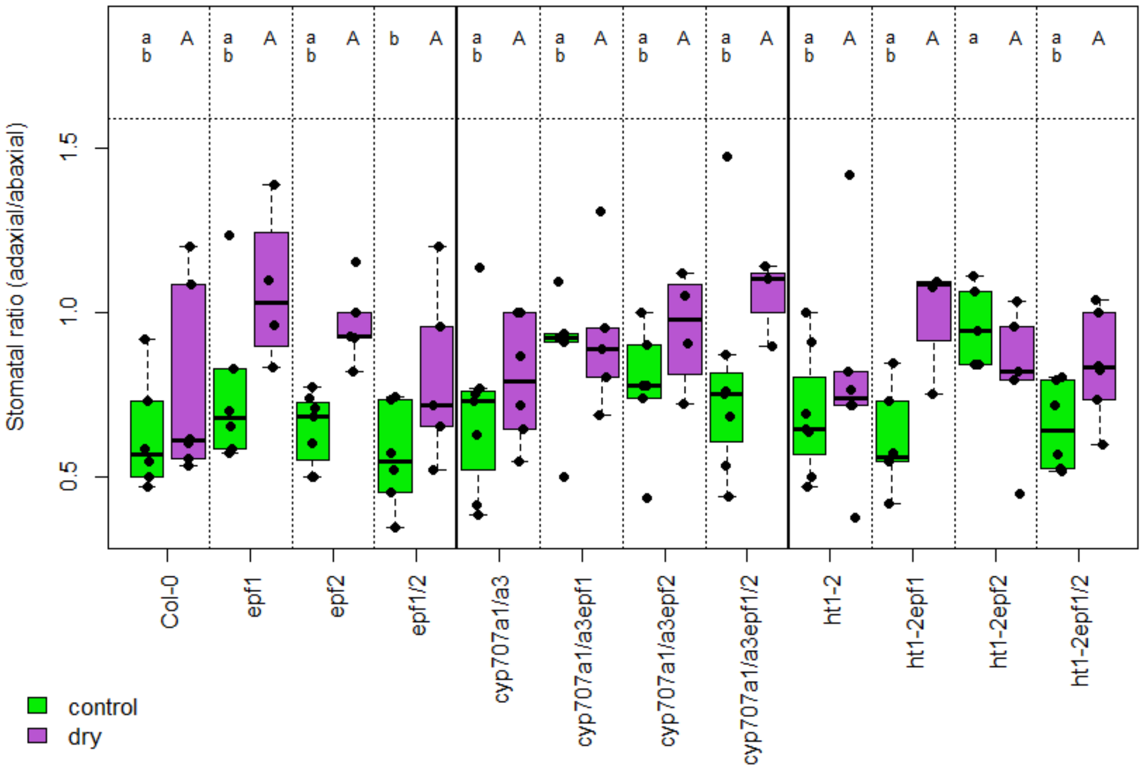


**Figure S4**. Stomatal ratios at the end of the growth experiment. Green boxes, normal growth RH; magenta boxes, low RH. The boxes span from the first to the third quartile, with median indicated with the horizontal line; the whiskers span the non-outlier range. Individual plants shown as solid dots. Asterisk above a pair of boxes indicates significant difference within that genotype according to Tukey's HSD (ANOVA involving all 24 treatment and genotype combinations). Shared lower case letters above the normal RH boxes indicate no significant difference according to Tukey HSD test (ANOVA involving only normal RH treatment); shared capital letters above the low RH boxes likewise for low RH treatment.

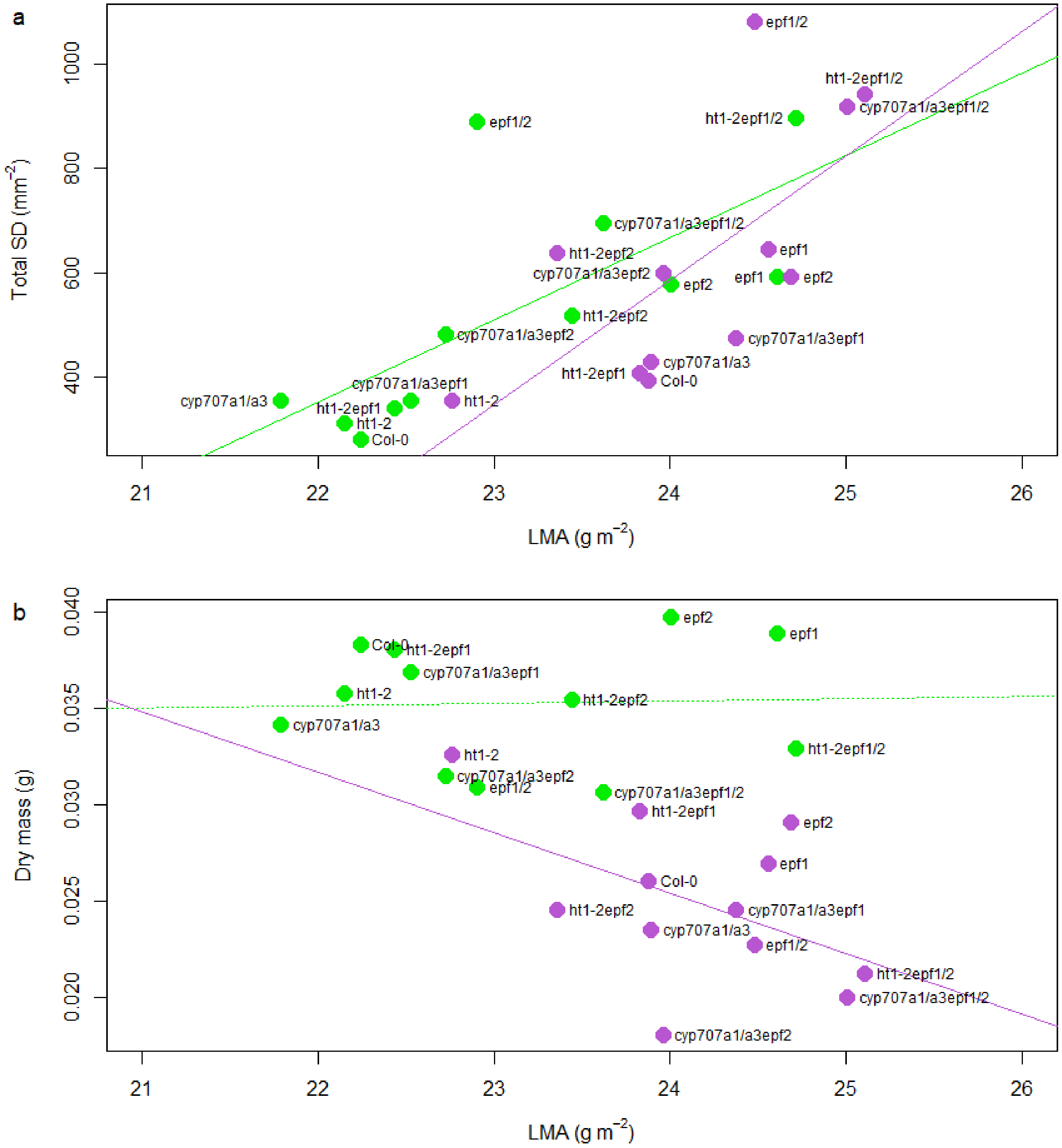


**Figure S5**. Relationship between (a) LMA and stomatal density, and (b) LMA and plant dry mass.Green markers, normal growth RH; magenta markers, low RH. Each dot indicates the median values (in both dimensions) of the shown plant line in given growth conditions. In (a), the slopes of the linear regression differ significantly from 0 at $P<0.05$, but not from each other. In (b), the linear regression between dry mass and LMA differs significantly from zero for the dry treatment plants.

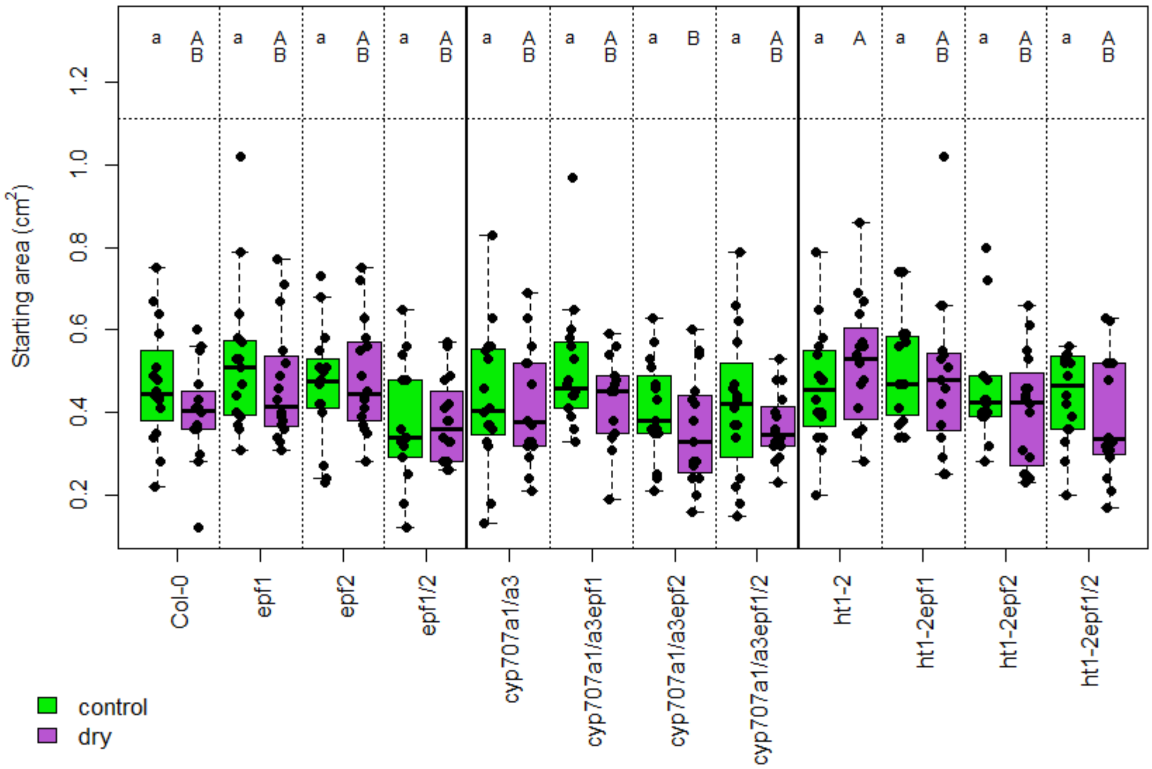


**Figure S6.** Projected areas of the growth experiment plants before the RH treatment started (day 14 after sowing). Green boxes, normal growth RH; magenta boxes, low RH. The boxes span from the first to the third quartile, with median indicated with the horizontal line; the whiskers span the non-outlier range. Individual plants shown as solid dots. Asterisk above a pair of boxes indicates significant difference within that genotype according to Tukey's HSD (ANOVA involving all 24 treatment and genotype combinations). Shared lower case letters above the normal RH boxes indicate no significant difference according to Tukey HSD test (ANOVA involving only normal RH treatment); shared capital letters above the low RH boxes likewise for low RH treatment.

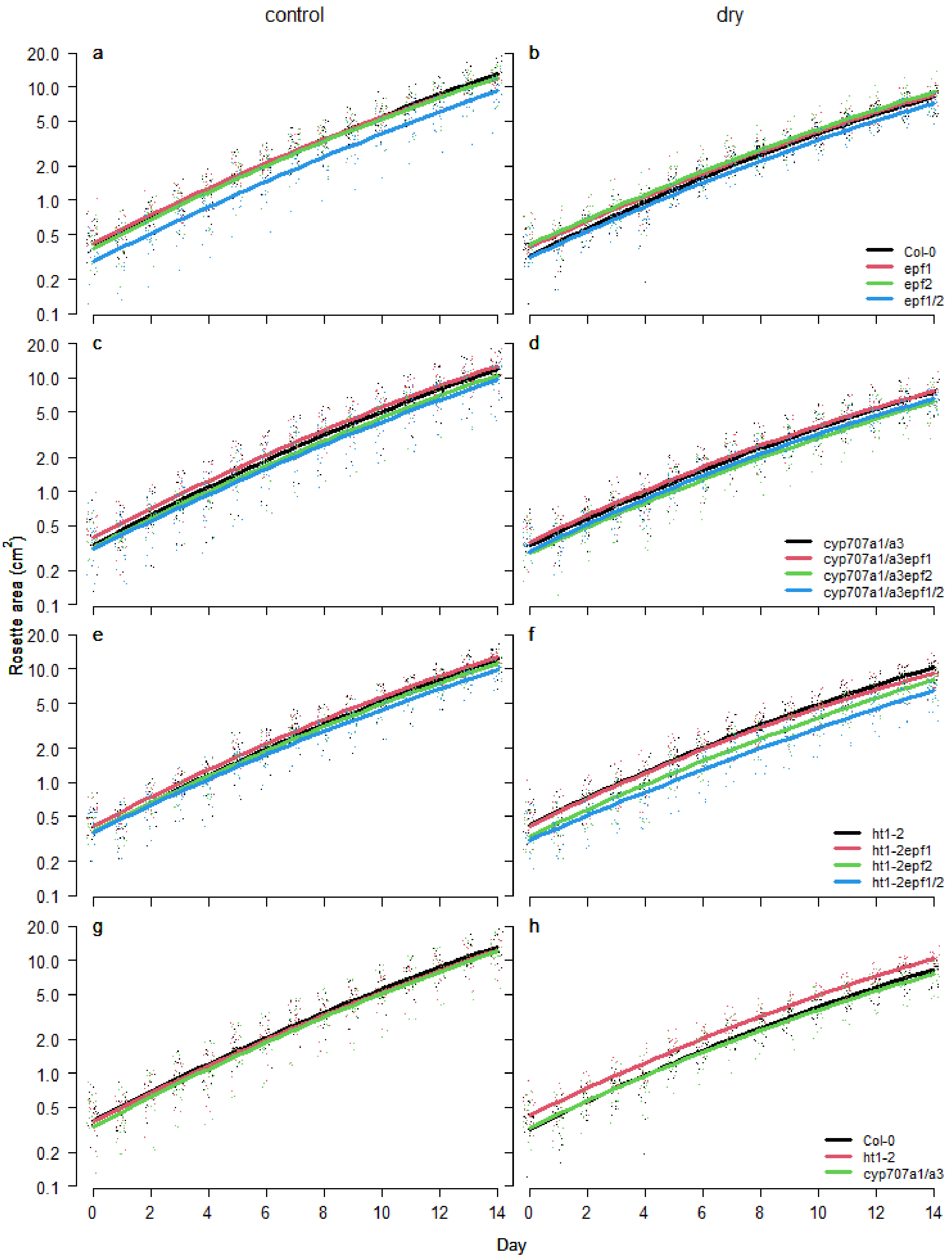


**Figure S7**. Time course of growth of the plant lines. (a,b), lines with no cyp707a1/a3 or ht1-2 mutations; (c,d), lines with *cyp707a1/a3* double mutation; (e,f), lines with *ht1-2* mutation; (g,h), lines with no *epf1* or *epf2* mutation. a,c,e,g, control conditions; b,d,f,h, low RH. Solid lines denote the mean of the ANCOVA (performed with log(area) as the dependent variable) estimate as in Table S3; y-axis values re-transformed to display raw projected area.
