## Supplementary Tables for "Low relative air humidity and increased stomatal density independently hamper growth in young Arabidopsis"

**Table S1.** Individual mutant lines used in this study and primers used for genotyping them.

| | | T-DNA border primer | Left primer (LP) | Right primer (RP) | Genotyping principle |
|---|---|---|---|---|---|
| *ht1-2* | Point mutation (Hashimoto et al. 2006) | | AGGATCCCAA CACACAAGGA | CATCTCGTCGTTCAA AAGCA | PCR product digested with PsuI (ThermoScientific) that cuts mutant but not wild-type product |
| *cyp707a1-1* | SALK_069127 | ATTTTGCCGATTT CGGAAC (LBb1.3) | CATGAACGTAT TGGGTTTTGG | TCCTGATATTGAATC CATCGC | Product with LP + RP from wild-type allele, and with T-DNA border primer + RP from mutant allele |
| *cyp707a3* | SALK_101566 | ATTTTGCCGATTT CGGAAC (LBb1.3) | GTTCCTGGAA GATTAATCGGC | ACGTGCTCTCGTCA CTCTCTC | Product with LP + RP from wild-type allele, and with T-DNA border primer + RP from mutant allele |
| *epf1-1* | SALK_137549 | ATTTTGCCGATTT CGGAAC (LBb1.3) | GGTGCATGTT CGACACTCTTC | CATGGTCATGTCCCG GAGAAGC | Product with LP + RP from wild-type allele, and with T-DNA border primer + RP from mutant allele |
| *epf2-2* | GABI_673E01 | ATATTGACCATCA TACTCATTGC | ATGACGAAGTT TGTACGCAAG | AATCTGATTCGGTCG TGTC | Product with LP + RP from wild-type allele, and with T-DNA border primer + RP from mutant allele |

**Table S2**. Pearson correlation coefficients between measured variables, with the median of each genotype at each growth RH forming one data point for each variable (12 levels for gas exchange data $g_l$, $A_n$, and WUE, 24 levels for others). Bold numbers indicate statistically significant correlation at $p<0.05$. SD, stomatal density on leaf 6; SC, stomatal count, SL, stomatal complex length; $g_l$, leaf conductance; $A_n$, net photosynthesis; WUE, intrinsic water use efficiency (the latter three at growth conditions); LMA, lea area/mass ratio; water content is percent mass loss during oven drying. Initial area is projected rosette area 14 days after sowing (at the beginning of the differential treatment); final area is area 28 days after sowing. Dry mass, LMA, and stomatal anatomy measured 29-30 days after sowing.

| | SD abaxial | SD adaxial | SD sum | SD ratio | SC abaxial | SC adaxial | SC sum | SL abaxial | SL adaxial | $g_l$ | $A_n$ | WUE | LMA | Water content | Initial area | Final area | Dry mass |
|---|---|---|---|---|---|---|---|---|---|---|---|---|---|---|---|---|---|
| SD abaxial | | **0.87** | **0.98** | -0.02 | **0.88** | **0.74** | **0.84** | -0.17 | -0.21 | **0.65** | **0.66** | **-0.69** | **0.57** | -0.33 | **-0.51** | **-0.50** | **-0.41** |
| SD adaxial | **0.87** | | **0.94** | 0.39 | **0.69** | **0.80** | **0.70** | **-0.47** | -0.36 | 0.49 | **0.67** | -0.56 | **0.75** | **-0.61** | **-0.58** | **-0.68** | **-0.62** |
| SD sum | **0.98** | **0.94** | | 0.12 | **0.85** | **0.81** | **0.84** | -0.30 | -0.31 | **0.61** | **0.65** | **-0.67** | **0.67** | **-0.43** | **-0.54** | **-0.58** | **-0.50** |
| SD ratio | -0.02 | 0.39 | 0.12 | | -0.18 | 0.24 | -0.09 | **-0.58** | -0.26 | -0.26 | 0.31 | 0.14 | **0.43** | **-0.65** | -0.18 | **-0.49** | **-0.49** |
| SC abaxial | **0.88** | **0.69** | **0.85** | -0.18 | | **0.85** | **0.97** | 0.06 | 0.00 | **0.73** | 0.53 | **-0.75** | **0.45** | -0.03 | -0.19 | -0.15 | -0.04 |
| SC adaxial | **0.74** | **0.80** | **0.81** | 0.24 | **0.85** | | **0.91** | -0.22 | -0.16 | **0.64** | 0.56 | **-0.69** | **0.68** | -0.29 | -0.16 | -0.29 | -0.17 |
| SC sum | **0.84** | **0.70** | **0.84** | -0.09 | **0.97** | **0.91** | | -0.03 | -0.09 | **0.70** | 0.44 | **-0.71** | **0.50** | -0.10 | -0.18 | -0.18 | -0.06 |
| SL abaxial | -0.17 | **-0.47** | -0.30 | **-0.58** | 0.06 | -0.22 | -0.03 | | **0.61** | 0.35 | -0.13 | -0.35 | **-0.51** | **0.78** | 0.33 | **0.75** | **0.76** |
| SL adaxial | -0.21 | -0.36 | -0.31 | -0.26 | 0.00 | -0.16 | -0.09 | **0.61** | | 0.30 | 0.00 | -0.16 | **-0.50** | **0.55** | 0.39 | **0.61** | **0.60** |
| $g_l$ | **0.65** | 0.49 | **0.61** | -0.26 | **0.73** | **0.64** | **0.70** | 0.35 | 0.30 | | 0.42 | **-0.93** | 0.17 | 0.11 | -0.42 | -0.10 | 0.02 |
| $A_n$ | **0.66** | **0.67** | **0.65** | 0.31 | 0.53 | 0.56 | 0.44 | -0.13 | 0.00 | 0.42 | | -0.39 | 0.56 | **-0.59** | -0.52 | **-0.63** | **-0.55** |
| WUE | **-0.69** | -0.56 | **-0.67** | 0.14 | **-0.75** | **-0.69** | **-0.71** | -0.35 | -0.16 | **-0.93** | -0.39 | | -0.30 | -0.06 | 0.47 | 0.16 | 0.05 |
| LMA | **0.57** | **0.75** | **0.67** | **0.43** | **0.45** | **0.68** | **0.50** | **-0.51** | **-0.50** | 0.17 | 0.56 | -0.30 | | **-0.68** | -0.22 | **-0.66** | **-0.55** |
| Water content | -0.33 | **-0.61** | **-0.43** | **-0.65** | -0.03 | -0.29 | -0.10 | **0.78** | **0.55** | 0.11 | **-0.59** | -0.06 | **-0.68** | | **0.43** | **0.92** | **0.90** |
| Initial area | **-0.51** | **-0.58** | **-0.54** | -0.18 | -0.19 | -0.16 | -0.18 | 0.33 | 0.39 | -0.42 | -0.52 | 0.47 | -0.22 | **0.43** | | **0.64** | **0.70** |
| Final area | **-0.50** | **-0.68** | **-0.58** | **-0.49** | -0.15 | -0.29 | -0.18 | **0.75** | **0.61** | -0.10 | **-0.63** | 0.16 | **-0.66** | **0.92** | **0.64** | | **0.97** |
| Dry mass | **-0.41** | **-0.62** | **-0.50** | **-0.49** | -0.04 | -0.17 | -0.06 | **0.76** | **0.60** | 0.02 | **-0.55** | 0.05 | **-0.55** | **0.90** | **0.70** | **0.97** | |

**Table S3.** Results of analysis of covariance to study the effect of growth RH on plant area. Dependent variable was log(projected area). Interactions of day$^2$ with other factors, as well as interactions of order 4 and higher were never significant and are not shown. Day*RH interaction indicates an overall gradual decline in plant size during the treatment; lack of interaction between day and genotype factors indicates that genotype-specific differences, where present, did not change during the treatment phase in weeks 3 and 4.

| | Sum Sq | Df | F value | Pr(>F) | |
|---|---|---|---|---|---|
| day | 637.4 | 1 | 7273.7809 | < 2.2e-16 | *** |
| day$^2$ | 26.8 | 1 | 306.1254 | < 2.2e-16 | *** |
| RH | 0.33 | 1 | 3.7669 | 0.0523269 | . |
| epf1 | 0.02 | 1 | 0.1809 | 0.6706260 | |
| epf2 | 3.72 | 1 | 42.4804 | 7.768e-11 | *** |
| sensitivity | 1.99 | 2 | 11.3486 | 1.206e-05 | *** |
| day*RH | 0.84 | 1 | 9.5698 | 0.0019879 | ** |
| day*epf1 | 0.03 | 1 | 0.3843 | 0.5353252 | |
| RH*epf1 | 0.00 | 1 | 0.0499 | 0.8231901 | |
| day*epf2 | 0.22 | 1 | 2.5409 | 0.1109886 | |
| RH*epf2 | 0.00 | 1 | 0.0009 | 0.9762573 | |
| epf1*epf2 | 2.20 | 1 | 25.1360 | 5.508e-07 | *** |
| day*sensitivity | 0.00 | 2 | 0.0006 | 0.9993646 | |
| RH*sensitivity | 0.06 | 2 | 0.3512 | 0.7038787 | |
| epf1*sensitivity | 0.52 | 2 | 2.9708 | 0.0513443 | . |
| epf2*sensitivity | 0.24 | 2 | 1.3468 | 0.2601642 | |
| day*RH*epf1 | 0.00 | 1 | 0.0283 | 0.8663185 | |
| day*RH*epf2 | 0.01 | 1 | 0.0651 | 0.7985699 | |
| day*epf1*epf2 | 0.00 | 1 | 0.0028 | 0.9574910 | |
| RH*epf1*epf2 | 0.03 | 1 | 0.3731 | 0.5413479 | |
| day*RH*sensitivity | 0.00 | 2 | 0.0226 | 0.9776270 | |
| day*epf1*sensitivity | 0.02 | 2 | 0.1197 | 0.8871570 | |
| RH*epf1*sensitivity | 0.27 | 2 | 1.5400 | 0.2144801 | |
| day*epf2*sensitivity | 0.01 | 2 | 0.0818 | 0.9214833 | |
| RH*epf2*sensitivity | 1.32 | 2 | 7.5058 | 0.0005555 | *** |
| epf1*epf2*sensitivity | 0.90 | 2 | 5.1246 | 0.0059769 | ** |
